# *Pseudomonas aeruginosa* differentiates substrate rigidity using retraction of type IV pili

**DOI:** 10.1101/2021.08.26.457786

**Authors:** Matthias D. Koch, Endao Han, Joshua W. Shaevitz, Zemer Gitai

## Abstract

The ability of eukaryotic cells to differentiate substrate stiffness is fundamental for many processes such as the development of stem cells into mature tissue. Here, we establish that bacteria feel their microenvironment in a similar manner. We show that *Pseudomonas aeruginosa* actively probes and measures substrate stiffness using type IV pili (TFP). The activity of the major virulence factor regulator Vfr is peaked with stiffness in a physiologically important range between 0.1 kPa (mucus) and 1000 kPa (cartilage). The local concentration of PilA at the base of dynamic TFP changes during extension and retraction in a surface dependent manner due to slow PilA diffusion in the cell membrane. Traction force measurements reveal that TFP retraction deforms even stiff substrates. Modeling of the measured substrate deformation and optical tweezers experiments suggest that TFP adhere at the tip only. Informed by these experimental results, we developed a model that describes substrate stiffness dependent dynamics of the polar PilA concentration which are quantitatively consistent with the transcriptional response to stiffness. Manipulating the ATPase activity of the TFP motors changes the TFP extension and retraction velocities and consequently the PilA concentration dynamics in a manner that is predictive of the experimental stiffness response. This work points to the use of a competition between PilA diffusion and TFP extension-retraction as a molecular shear rheometer. Our results highlight that stiffness sensing is a conserved property between the kingdoms of life.

## Introduction

Bacteria thrive in many physically different environments and have developed strategies to discern among using physical cues [1]. In many bacteria, surface attachment promotes the formation of robust biofilms that offer protection from environmental insults such as fluid flow or harmful chemical exposure such as antibiotics [2]. The transition from planktonic to surface association triggers genetic programs that promote surface attachment and biofilm formation [3]. Consequently, the ability to sense a surface is crucial for bacteria during this process. Surface sensing can be facilitated passively by membrane associated proteins that sense the presence of a surface upon contact or actively by obstruction of the motion of an extracellular appendage [4–9]. For example, flagella are long helical appendages that generate a propulsive force that enable bacteria to swim in liquid. Attachment of either the cell body or the flagellum to the surface changes the torque on the motor which results in stator remodeling and consequent downstream signaling [10–12]. In contrast, bacterial type IV pili (TFP) are mostly straight filamentous appendages with a length of about one micron that drive surface bound twitching motility through cycles of extension and retraction [13]. *Caulobacter crescentus* has been shown to detect the presence of a surface when TFP retraction stalls as cells adhere to the surface [14]. This rapidly triggers the formation of a highly sticky stalk that promotes surface attachment.

In both examples above, the surface is detected in a binary fashion, the cell is either near a surface or not. While detecting the presence of a substrate is a remarkable feature on its own, different solids can vary in their mechanical properties tremendously and can be as rigid as bone or as soft as mucus. Eukaryotic cells integrate a much larger variety of mechanical cues and can differentiate substrates by stiffness [15, 16]. Stem cells, for example, differentiate into different types of tissue depending on substrate stiffness and malfunctioning of this mechanotransduction process can result in severe diseases [17, 18].

Stiffness is the ratio of force to deformation. Sensing substrate stiffness requires a cell to actively deform the substrate, measure the resulting force-deformation relationship, and transduce this information into a biochemical signal. A common pathway in eukaryotic cells uses contraction mediated by focal adhesions, molecular motors, and the actin cytoskeleton. These assemblies can generate substantial forces in the nanonewton range. Information about the substrate stiffness is sensed by conformational changes in the integrin complexes and processed by phosphorylation of focal adhesion kinases [19]. Bacteria are much smaller and conversely it is harder for them to generate a large force to deform a substrate and probe its mechanical properties [20].

In the human body, commensal and pathogenic bacteria encounter a broad spectrum of mechanically different surfaces ranging from slimy mucus in the lung to soft tissue of wounds to stiff bones. *Pseudomonas aeruginosa*, for example, is an opportunistic human pathogen that infects all of these mechanically different sites [21]. A crucial determinant of *P. aeruginosa* virulence are TFP and TFP-mediated surface sensing has been implicated in multiple virulence pathways [9, 22–24]. We previously showed that TFP retraction on hydrogels stimulates the production of cAMP which activates the virulence factor regulator (Vfr), a major transcriptional regulator for over 100 virulence related genes including the PaQa operon [25]. In these experiments, Vfr activity was monitored fluorescently. We engineered the transcriptional fluorescent reporter P_PaQa_::YFP such that YFP fluorescence increases with Vfr activity. We found that Vfr was upregulated on diverse hydrogels compared to its activity in planktonic cells. Thus, P_PaQa_::YFP fluorescence is a direct readout of the substrate dependent transcriptional activity of Vfr.

The P_PaQa_ response increased on gels with increasing agarose concentration in the range of 0.75% to 1.5% agarose. This increase could be due to an increase in substrate stiffness, indicative of stiffness sensing, or due to a decrease in hydrogel pore size, indicative of a sensory system that responds to surface topography. Given that TFP are powered by a strong molecular motor able to generate more than 150 piconewton of force, we here investigate the hypothesis that TFP retraction facilitates substrate stiffness sensing [26, 27]. We show that *P. aeruginosa* responds to substrate stiffness and that this response is peaked, i.e. nonmonotonic. We show that TFP retraction can probe and deform the substrate and that the local concentration of PilA at the TFP base changes during TFP extension-retraction in a substrate dependent manner. Based on these results, we propose a quantitative model for substrate stiffness sensing that measures temporal changes in the local PilA concentration during TFP activity on different substrates. This model is consistent with stiffness sensing results of different mutants affecting PilA dynamics.

## Results

### cAMP production depends on substrate stiffness and is peaked at intermediate rigidities

To investigate if the substrate and cAMP-dependent response of P_PaQa_ is a result of increasing stiffness or decreasing pore size for agarose gels with increasing concentration, we conceived an assay that utilizes two chemically different stiffness-tunable hydrogels: polyacrylamide (PAA) and agarose. By changing the concentration of the polymer acrylamide and its crosslinker bis-acrylamide, the stiffness of PAA gels can be adjusted between 0.1 kPa and 500 kPa [28]. Similarly, by adjusting the concentration of agarose, the stiffness of agarose gels can be adjusted between 1 kPa and 1000 kPa [29]. As shown in Fig. 1A and explained in the Supplementary Materials and Methods section, the pore size of both gel types differs by about one order of magnitude for any given stiffness [30–33]. This difference in pore size and stiffness allows us to disentangle a cellular response to either of these physical gel properties. We measured the P_PaQa_ response to a variety of these hydrogels and plotted the results separately as a function of pore size and a function of stiffness. If the P_PaQa_ response is a result of the changes in pore size, then we expect the P_PaQa_ response data for agarose and PAA to collapse onto each other as a function of pore size, but not as a function of stiffness. In contrast, if the P_PaQa_ response is a result of a change in stiffness, then the response for agarose and PAA should collapse as a function of stiffness, but not as a function of pore size.

**Fig. 1.**
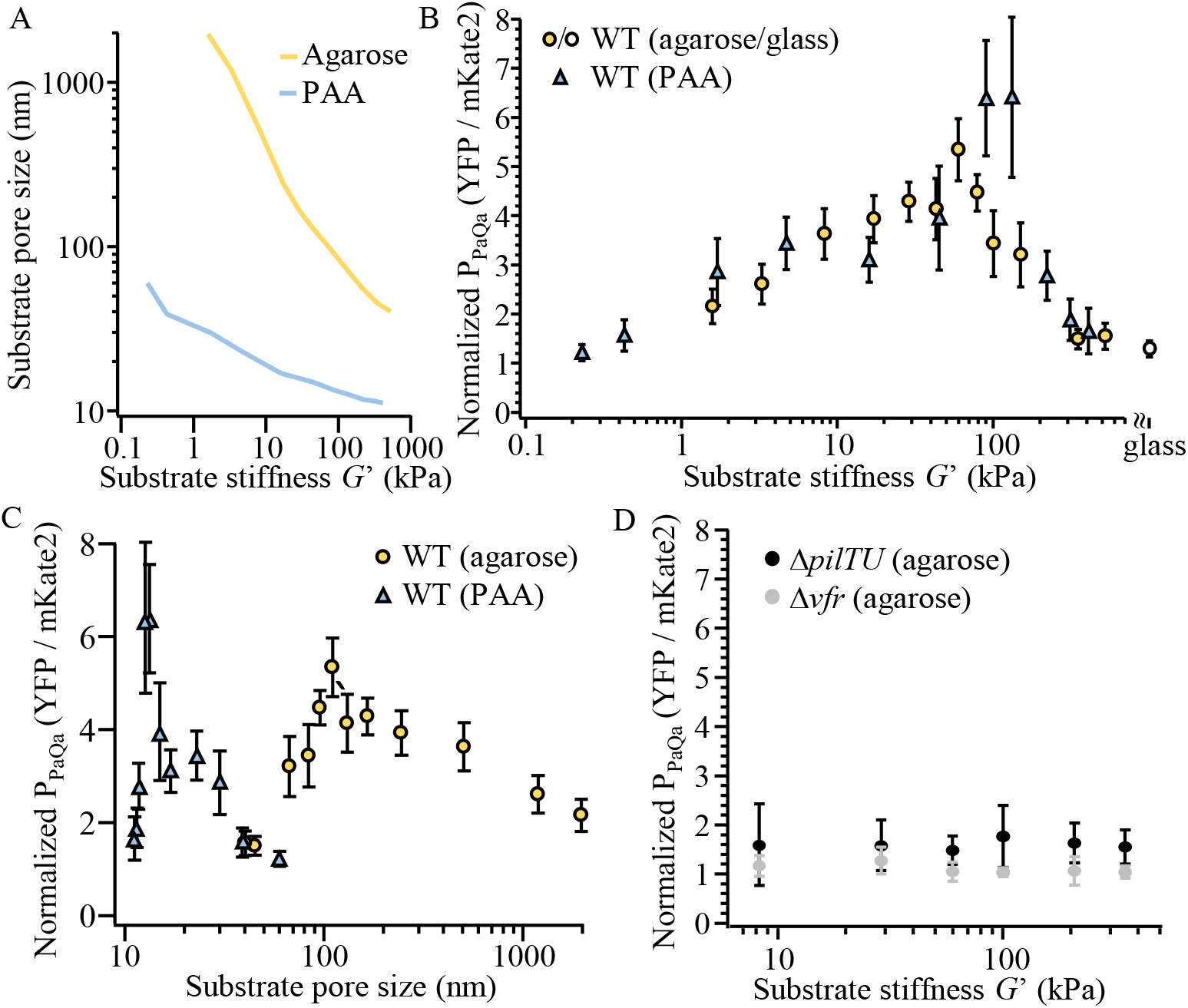
*P. aeruginosa* senses substrate stiffness and the transcriptional stiffness response is peaked at intermediate stiffness. (A) The pore size of agarose and PAA hydrogels differs by about one order of magnitude for each stiffness. The data are compiled from multiple publications (see Results and Materials and Methods). (B) The maximum expression of P_PaQa_ for cells on agarose and PAA hydrogels and on a glass coverslip normalized to the expression in liquid collapses as a function of substrate stiffness and is peaked between 50 – 100 kPa. (C) The response of P_PaQa_ on agarose and PAA hydrogels does not collapse as a function of substrate pore size. (D) The P_PaQa_ responses of the retraction deficient mutant Δ*pilTU* and the mutant lacking the transcriptional regulator Δ*vfr* do not respond to stiffness. (B-D) Each data point is the median of at least three biological replicates and 8 technical replicates each. Error bars are the inter quantile range (IQR) of the distribution (See SI Fig. 1).

To measure the cAMP dependent response to these substrates, we recorded individual fluorescence timelapse movies of *P. aeruginosa* cells growing on a variety of different PAA and agarose gels. We then analyzed the P_PaQa_ response to a variety of these gels. Specifically, we analyzed the increase in P_PaQa_ fluorescence compared to the constitutive fluorescence of P_rpoD_ over time, where the zero-time point represents the P_PaQa_ expression in liquid. The expression of P_PaQa_::YFP is maximal after about three hours (Supplementary Fig. 1A-C). Consequently, we analyzed the median and inter quantile range (IQR) of the distributions of relative fluorescence values P_PaQa_ / P_rpoD_ (YFP / mKate2) at *t* = 0 (liquid) and the maximum response for each technical repeat around *t* = 3 hrs. The ratios of these medians represent the upregulation as a response to the hydrogel compared to the expression in liquid (Supplementary Fig. 1D).

We first compared the P_PaQa_ response on PAA to that on agarose as a function of substrate stiffness (Fig. 1B). The measured P_PaQa_ responses for both gel types collapse almost perfectly onto each other over a four orders of magnitude range in stiffness, suggesting that the P_PaQa_ response to a hydrogel is a result of a change in stiffness. Interestingly, the P_PaQa_ response is not monotonically increasing with increasing stiffness, but is peaked at an intermediate stiffness of 50 – 100 kPa and decreases for stiffer gels. To test if this high stiffness decrease is a possible hidden effect of ever decreasing pore sizes, we next plotted the same P_PaQa_ response data as a function of pore size (Fig. 1C). Here, the responses for both gel types do not collapse onto each other. In fact, both responses do not overlap at all and the response on PAA peaks below the smallest pore size of all agarose gels supporting that the observed high stiffness decrease of P_PaQa_ is not a hidden effect of a small pore size but rather a response to high substrate stiffness. In contrast, the doubling time of individual cells during the P_PaQa_ timelapse experiments depends on pore size but not stiffness (Supplementary Fig. 2). We conclude that the substrate dependent P_PaQa_ response is a response to stiffness and that this response is peaked at intermediate substrate stiffness.

We previously demonstrated that the P_PaQa_ substrate response depends on TFP retraction, and we sought to check if this is also true for stiffness sensing. As shown in Fig. 1B, the P_PaQa_ response of a Δ*pilTU* mutant that still makes TFP but is unable to retract the TFP is independent of surface stiffness. Similarly, disrupting the sensing pathway by deleting the virulence response regulator Vfr also abolishes stiffness sensing.

### The local concentration of PilA at the base of TFP changes during TFP extension-retraction in a surface dependent manner

The P_PaQa_ response to stiffness relies on retraction of TFP. Interestingly, we observed that the local concentration of PilA at the base of dynamic TFP changes during TFP extension and retraction. We fluorescently labeled PilA employing a cysteine-maleimide labeling technique [34] and measured the local PilA fluorescence intensity at the base of extending TFP that are not in surface contact (Fig. 2A,B). The pilus shown in Figure 2A extended for about 13 s to a length of about 5 μm. During this time, the fluorescence intensity at the pilus base decreased by about 20%. Immediately after, the pilus retracted completely back into the cell for about 7 s and the fluorescence intensity at the base increased simultaneously. Interestingly, upon retraction the intensity initially increased to higher levels than before pilus extension, only to decreases slowly again to initial levels several seconds after the pilus was fully retracted. This shows that the concentration of PilA at the base of dynamic pilus changes significantly during extension and retraction. We wondered if this behavior changes for TFP attached to a stiff substrate as sensing PilA substrate dependent concentration changes might be a mechanism for stiffness sensing. We thus compared the above analysis of the local PilA concentration for unbound TFP to the change of PilA concentration of TFP bound to a stiff 4.0% (350 kPa) agarose gel (Fig. 2C,D). Here, the pilus extended to 3 μm length and retraction stalled immediately as the tip adhered to the gel, indicated by a straining of the pilus and a slight movement of the cell towards the TFP tip (Supplementary Movie 6). As expected, the fluorescence intensity at the pilus base increased slowly upon stalling but did not overshoot the initial intensity level compared to unbound pili (Fig. 2A,B). Once equilibrated by diffusion (*t* > 15 s), the fluorescence level remained lower than the initial level prior to pilus extension because a fraction of PilA remained outside of the cell in the extended pilus. Thus, dynamic changes of the PilA concentration at the pilus base change with the degree to which TFP retract, which in turn depends on the substrate stiffness.

**Fig. 2.**
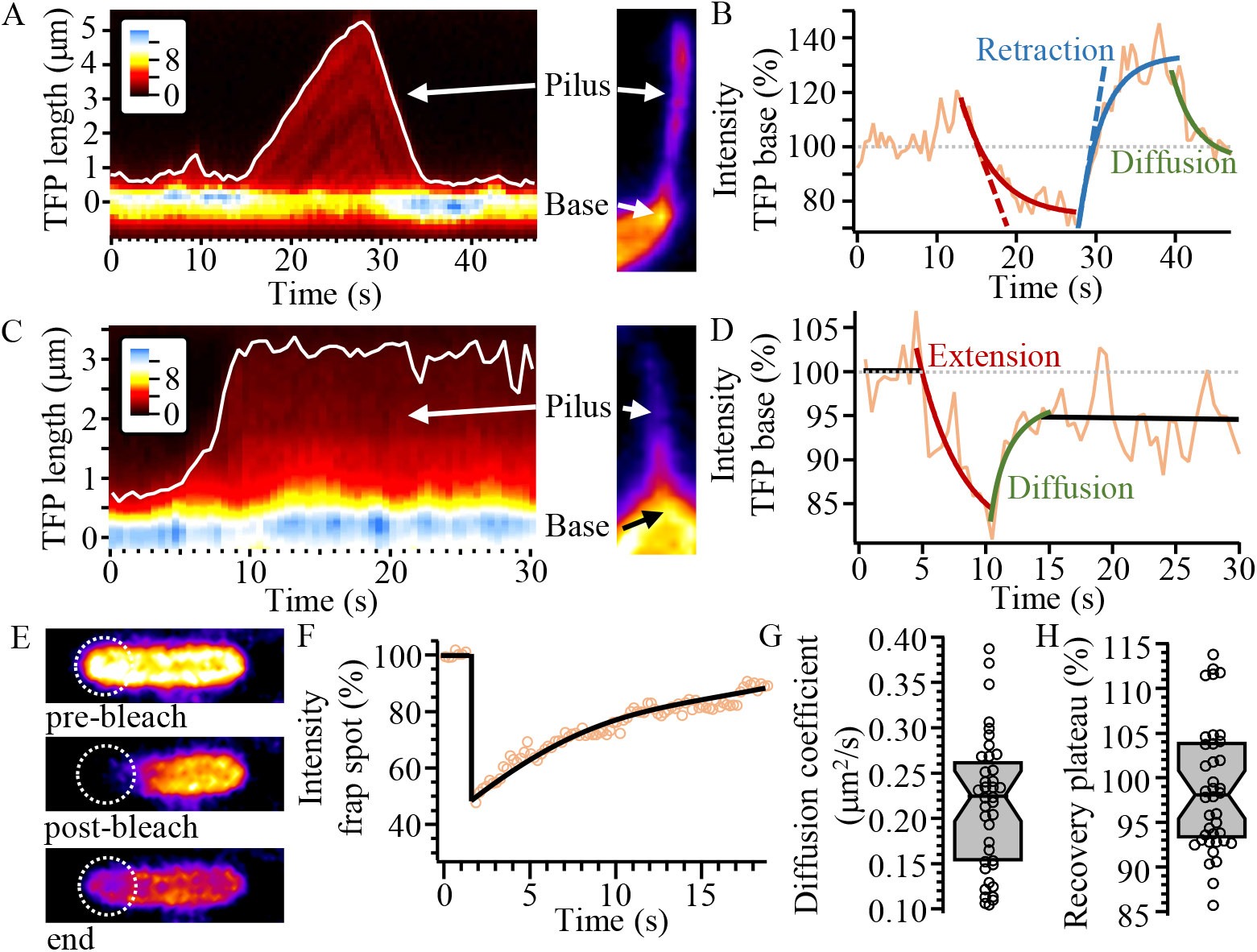
The local concentration of PilA at the base of dynamics TFP changes during TFP extension-retraction in a substrate dependent manner. (A) Kymograph of the PilA-Alexa488 fluorescence intensity along the dynamic pilus without surface attachment shown on the right (Inset: fluorescence intensity (a.u.)). (B) Analysis of the fluorescence intensity from A) at the pilus base indicative of the concentration of PilA. Thick/dashed red, blue, and green lines are exponential/linear fits. (C) Kymograph of the PilA-Alexa488 fluorescence intensity along the dynamic pilus attached to a 4.0 % agarose gel shown on the right (Inset: fluorescence intensity (a.u.)). (D) Analysis of the fluorescence intensity from C) at the TFP base indicative of the concentration of PilA. Thick red and green lines are exponential fits. (E) Three frames of a fluorescence recovery after photobleaching (FRAP) experiment of Alexa488 labeled PilA. The dashed circle illustrates the photobleaching spot. (F) Fluorescence intensity at the photobleaching spot in E) over time, corrected for overall bleaching (see Supplementary Methods). The thick black line for t > 1.5 s is an exponential fit. (G) Diffusion coefficient obtained by individual FRAP experiments. (N = 39). (H) Recovery plateau of post bleach fluorescence at the bleach spot compared to before bleaching, indicative of the immobile fraction of PilA, obtained by individual FRAP experiments (N = 39). G,H) Gray boxes are the median and 25/75% quantiles.

### Diffusion of PilA in the inner membrane is slow compared to TFP extension and retraction

Surprisingly, the result that the local PilA concentration changes during TFP extension/retraction also suggests that the diffusion of individual PilA proteins in the inner membrane is slower than the dynamics of TFP themselves. This can fundamentally limit TFP function since the TFP assembly complex could locally run out of monomers to assemble into extending TFP long before the entire cellular pilin subunit pool is depleted. To test this idea, we used fluorescence recovery after photobleaching (FRAP) to measure the diffusion coefficient of PilA in the inner membrane. We photobleached PilA fluorescence at one pole of the cell with a focused laser for less than 1 second and measured the dynamics of fluorescence recovery due to diffusion of unbleached monomers from the rest of the cell (Supplementary Methods, Fig. 2E,F) [35]. This analysis reveals a diffusion coefficient of *D*_PilA_ = 0.22 μm^2^/s which is comparable to, but slower than, the diffusion of other inner membrane proteins in *E. Coli* (Fig. 2G and Supplementary Methods) [36]. We also found that the fluorescence intensity at the bleached pole recovers to the initial level (corrected for overall photobleaching), indicating that there is no immobile fraction of PilA monomers that are not diffusing and locally stuck (Fig. 2H).

The loss of PilA during TFP extension can be compared to the effect of diffusion by calculating a corresponding diffusion coefficient for PilA extension: a typical pilus with length *L* = 1 μm and extension velocity *v*_ext_ = 350 nm/s can be described by an effective diffusion coefficient *D*_ext_ = *v*_ext_ · *L* = 0.35 μm^2^/s. Even more dramatically, the pilus shown in Figure 2A, with length 5 μm, yields an effective diffusion coefficient of *D*_ext_ = 1.5 μm^2^/s. This shows that the loss of PilA due to extension of a pilus dominates over the diffusive behavior of PilA in the membrane in governing the concentration of PilA at the pilus base. The slow diffusion of pilins in the membrane might be one reason that PilA is made in high abundance in the cell [37].

### Forceful pilus retraction deforms even rigid substrates

How can individual cells sense the stiffness of their surrounding substrate? Stiffness is the degree to which a material resists deformation. The stiffness of a spring for example is given by the spring constant *k* = *F*/Δ*x* and is a measure for the spring’s resistance to an applied force *F* that yields a stretch Δ*x*. Similarly, the one-dimensional deformation Δ*x* of the surface layer of a gel with the shear modulus *G*’ by the force *F* is given by 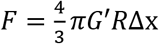, where *R* is the radius of the spot the force is applied to (see Supplementary Methods and Supplementary Fig. 3A). In other words, any stiffness sensing mechanism must be able to apply a force and measure the resulting deformation. Since the stiffness-dependent P_PaQa_ response relies on TFP retraction, we thus hypothesized that substrate stiffness could be sensed by a molecular sensor that measures the force and deformation of the substrate when gel-attached TFP retract. To test this hypothesis, we first determined the mechanical properties of PAO1 TFP retraction with an optical trap (Fig. 3A) as the retraction force of the TFP of different strains varies significantly and the rate at which TFP slow down under load has only been reported for the bacterium *Neisseria gonorrhoeae* [26, 27, 38–41]. Clean 500 nm diameter polystyrene beads are trapped with a highly focused laser beam and brought in proximity of a cell pole. TFP readily bind to polystyrene and retraction of bead bound TFP can be observed by a displacement of the bead towards the cells pole [39]. The optically trapped bead initially gets displaced by pilus retraction with the load free velocity *v*_0_, and then slows down due to an increase in retraction force, until the bead stalls (Fig. 3B). The pilus stall force or maximum force a single *P. aeruginosa* PAO1 pilus retraction motor can generate was measured to be *F_s_* = 55 pN (Fig. 2C). This value agrees to the ranges of TFP retraction forces measured in other species such *P. aeruginosa* PA14, *Myxococcus*, or *Neisseria* [27, 39, 41]. We also measured the load free TFP retraction velocity to be *v*_0_ = 600 nm/s, in good agreement with observations by fluorescence microscopy (Fig. 2D) [34, 42]. Further, we determined the force-velocity relationship that describes the decrease in retraction velocity *v*(*F*) = *v*_0_ – *f*_0_ · *F*, with *f*_0_ = −13 nm/(s·pN) (Fig. 2E, Supplementary Fig. 3). This parameter will become important later in this manuscript when we present a model for stiffness sensing. These results group the TFP of PAO1 among the most forceful TFP, a prerequisite to significantly deform a surface.

**Fig. 3.**
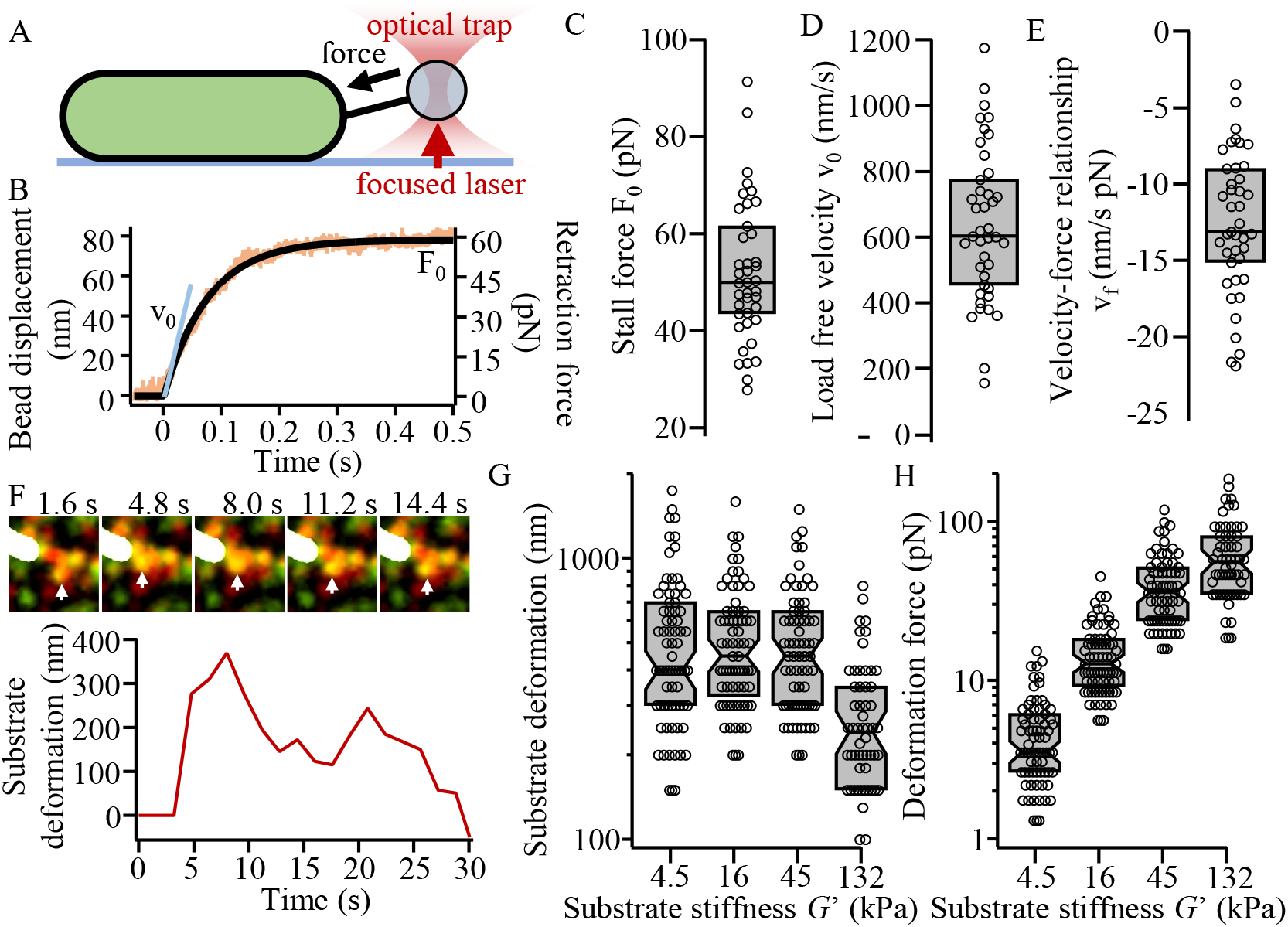
Forceful TFP retraction deforms and force probes stiff substrates. (A) Sketch of optical trapping experiment. An optically trapped bead is brought close to the pole of a coverslip attached cell. The bead is displaced towards the cell upon TFP binding and retraction. (B) Exemplary bead displacement/pilus retraction trajectory. The TFP retracted with initial velocity *v*_0_ ≈ 600 nm/s, then slowed down due to increasing resistance or retraction force from the trap until it stalls with the maximum stall force *F*_0_ ≈ 60 pN. (C) Distribution of retraction stall forces of individual pili (N = 56). (D) Distribution of load free velocities of individual TFP retractions (N = 56). (E) Distribution of the velocity-force relationship of individual pilus retractions (N = 56). See Supplementary Fig. 3 and Supplementary Methods for more details. (F) TFM example of a substrate deformation (white arrow) caused by pilus retraction towards the cell on a stiff 132 kPa PAA gel. The substrate deformation was obtained by particle tracking. (See Supplementary Movie 5). (G) Substrate deformation of individual pilus retraction events on substrates with different stiffness obtained by particle tracking (N > 58 each). (H) Deformation force for the individual deformations in G) as a function of substrate stiffness (N > 56 each). C-H) Gray boxes are the median and 25/75% quantiles.

To further test the hypothesis that *P. aeruginosa* deforms and probes substrates using force, we next used traction force microscopy (TFM) to see if TFP retraction causes local substrate deformation. In TFM, small 40 nm diameter fluorescence beads are embedded densely into the hydrogel substrate and imaged in a timelapse experiment. Substrate deformations become apparent because gel-bound beads get displaced together with the substrate (Fig. 3F). TFP caused substrate deformations in front of the pole of individual cells on both soft 4.5 kPa gels and stiff 132 kPa gels (Supplementary Movies 1-4). We used single particle tracking to measure the surface deformation caused by individual cells for the different substrates (Supplementary Methods). This analysis shows that even for the stiff 132 kPa substrates, TFP are still able to cause measurable deformation (Fig. 3F, Supplementary Movie 5). In the example shown here, pilus retraction stalled slowly over 4 seconds and deformed the gel by approximately 350 nm before slowly relaxing back (Fig. 3F). Quantifying substrate deformations for a range of gel stiffnesses shows that gel deformation is maximal for soft substrates and declines only for very stiff substrates (Fig. 3G). The fact that substrate deformations are constant for substrate stiffnesses < 100 kPA suggests that deformations are limited by TFP length, because TFP fully retract. Only on stiff gels surface deformations are not limited by TFP length but TFP retraction force. Consequently, the deformation force for stiff 132 kPa gels equals the maximum TFP stall force obtained by the optical trapping experiments above. Thus, setting the median deformation force equal to the stall force we obtain the TFP attachment spot size 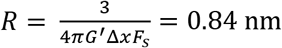. Scaling the substrate deformaions this ways shows that the traction force applied to deform the substrate increases monotonically with substrate stiffness (Fig. 3H).. Together, these results show that TFP can deform even stiff substrates in support of the hypothesis that TFP retraction causes substrate deformation, which can be sensed by a molecular stiffness sensor.

### Temporal changes of the local pilin concentration yield a stiffness dependent input signal for a PilA sensory system

Our results show that TFP retraction deforms hydrogel substrates, that the resulting deformation and the applied traction force depends on the stiffness of the substrate, that the concentration of PilA at the base of TFP is dynamic, and that these dynamics change with substrate stiffness. Together, these results support a model where the retraction of gel-bound TFP deforms the substrate and information of the substrate dependent force-deformation relation (i.e., substrate stiffness) is read out by a cell via substrate dependent changes in the dynamics of the local PilA concentration at the base of TFP (Fig. 4A). We used a two-tiered modeling approach to simulate changes in PilA concentration during TFP retraction for different substrate stiffnesses.

**Fig. 4.**
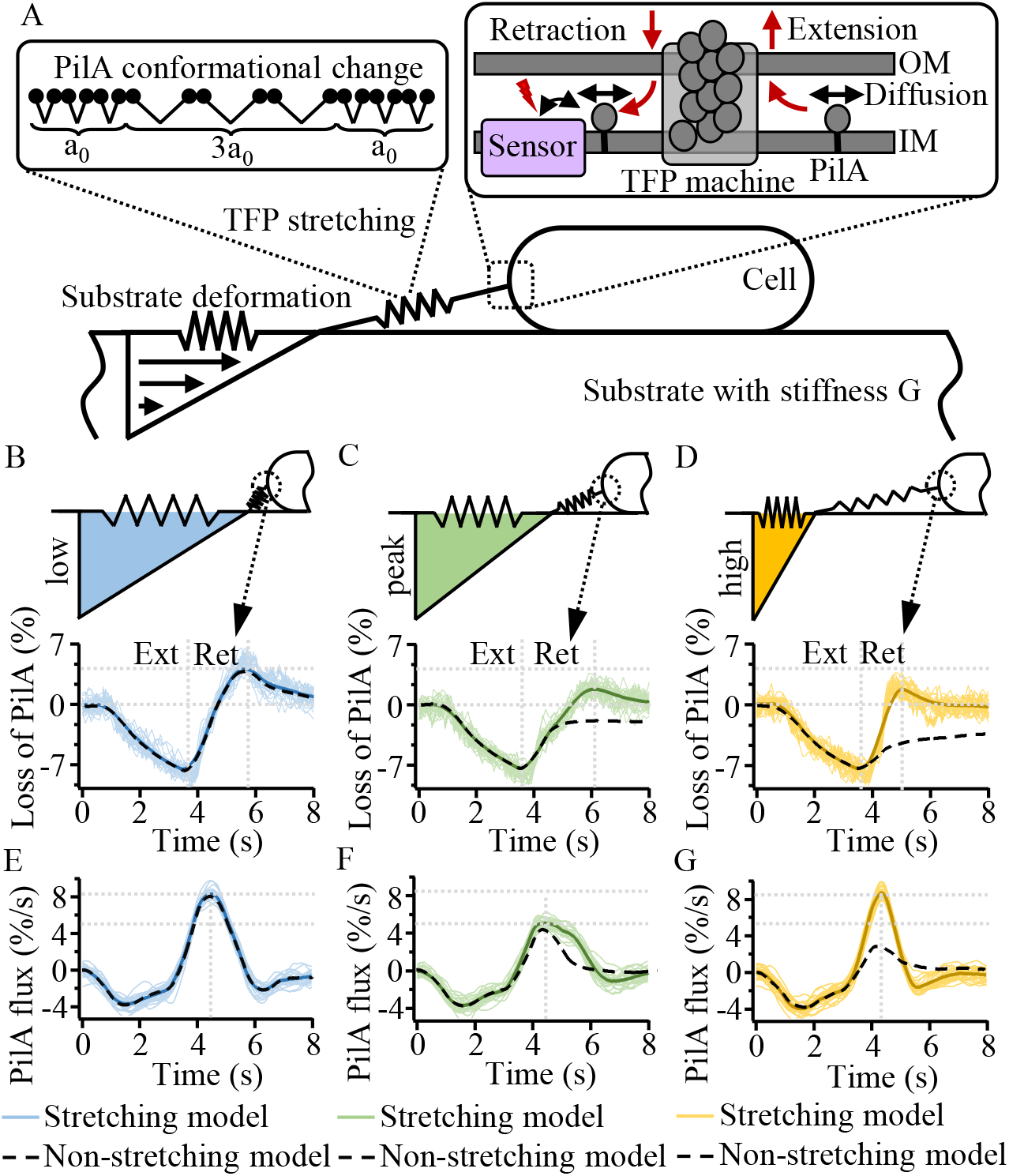
Model and simulation of the TFP retraction dependent changes in the local PilA concentration at the base of TFP on different substrates. (A) Sketch of the model. TFP retraction leads to elastic substrate deformation. The TFP fiber is modeled as stiff and stretching fiber, where stretching is facilitated by conformational changes of PilA (top left inset). Extension and retraction result in local changes of the Pila concentration at the pole that can be sensed by a molecular PilA sensor (top right inset). (B) - (D) The local PilA concentration at the base of TFP during TFP extension (Ext) and TFP retraction (Ret) for a gel with B) low, C) intermediate (at the peak of the PaQa response), and D) high stiffness. (E) - (G) Temporal change or flux of the local PilA concentration at the base of TFP during TFP extension (Ext) and TFP retraction (Ret) for a gel with E) low, F) intermediate (at the peak of the PaQa response), and G) high stiffness. (B) - (G) Black dashed lines are the median of 50 individual simulations of the non-stretching TFP model. Bold colored lines are the median of 50 individual simulations of the TFP stretching model, each individual simulation is shown as thin colored line.

First, we combined the experimental TFM and optical trapping results to describe the mechanical coupling of the retraction of substrate-attached TFP and the substrate deformation in a numerical simulation (Supplementary Fig. 4). In addition to the effect of force on retraction speed we measured using the optical trap, TFP also exhibit a force-induced reversible stretching under extensile loads likely due to a conformational change in the PilA monomers. This unfolding like feature was first demonstrated in gonococcal TFP but has also been shown for TFP in *Vibrio cholerae* and *P. aeruginosa* [38, 43–46]. As shown in Supplementary Fig. 4, the TFP retraction force that deforms the substrate ramps up more quickly on stiff gels compared to soft gels. Consequently, the tension in PilA monomers increases more rapidly on stiff gels. PilA thus gets stretched more quickly on stiff gels which increases the ratio of stretched to unstretched monomers in the pilus (Supplementary Fig. 4E). More unstretched monomer increases the pilus rest length and thus more monomers that can be retracked back into the cell compared to non-stretching TFP. In fact, looking at the loss of PilA from the inner membrane at a fixed time point after the initiation of TFP retraction reveals a profile that is peaked with stiffness. We empirically determined that a fixed time point of 1 s is ideal to fit the model response to the experimental P_PaQa_ data and determine optimal values for the free fit parameters (the equilibrium folding and unfolding rates of individual monomers and the attachment spot size of the TFP to the substrate). The resulting model explains the experimental P_PaQa_ response reported above (Supplementary Fig. 4H).

Encouraged by these results, we coupled the numerical mechanics simulation that describes the simulated influx of PilA over time to a simulation of stochastic PilA diffusion simulation in the inner membrane (Fig. 4A). This combined simulation is necessary to adequately describe the competition on different time scales between PilA diffusion and the TFP extension-retraction dependent outflux/influx of PilA. As shown in Fig. 4B-D, TFP extension yields a substrate stiffness independent loss of the concentration of PilA in the inner membrane. The simulated influx of PilA during TFP retraction on a soft gel yields an increase in the local PilA concentration above the level before TFP extension started, in good agreement to our experimental observation (Fig. 4B, colored lines).

We analyzed the temporal changes of the local PilA concentration during extension and retraction on different stiffnesses (Fig. 4E-G). We find that the influx of PilA is maximal for soft and stiff gels, and lower for intermediate stiffnesses. Plotting the maximum influx of PilA during TFP retraction as a function of substrate stiffness reveals that the change in the local PilA concentration is peaked at intermediate stiffness for the stretching model and quantitatively explains the experimentally observed PaQa stiffness response (Fig. 5A). This result suggests that the dynamic changes of PilA concentration that we also observed experimentally (Fig. 2) can yield a stiffness dependent input signal for a PilA sensory system. Interestingly, if the PilA subunits are not allowed to stretch, the maximum influx of PilA steadily decreases with substrate stiffness and does not show a peak at intermediate stiffnesses (Fig. 4B, black lines). This suggests that stretching of PilA might be a key aspect of the mechanism that yields a peaked stiffness response.

**Fig. 5.**
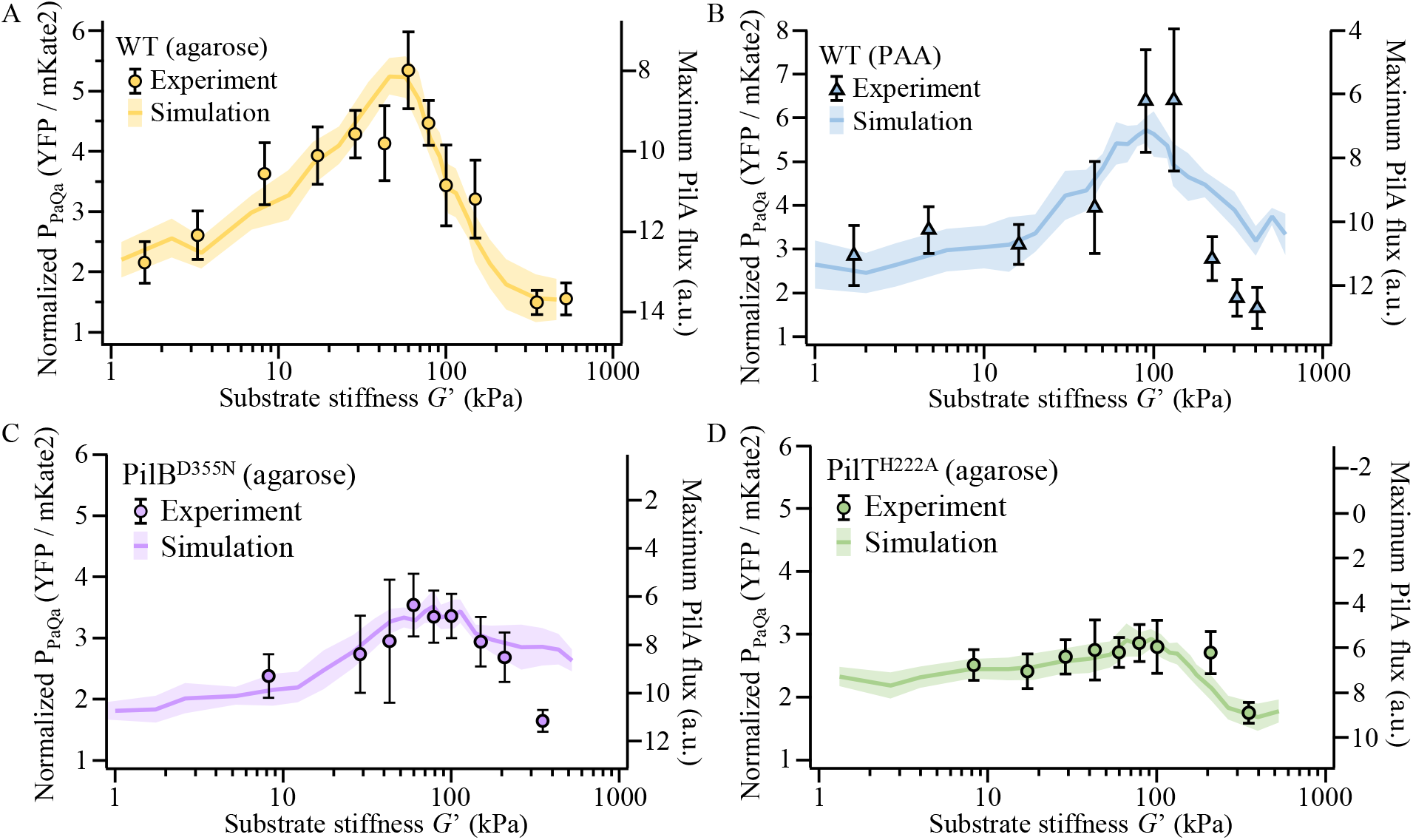
Changes in the TFP extension or retraction velocity change the response to substrate stiffness in a manner that is predictive by the model. (A) *P. aeruginosa* PAO1 WT cells on agarose. (B) *P. aeruginosa* PAO1 WT cells on PAA. (C) The slowly extending ATPase point mutant PilB^D355N^ on agarose. (D) The slowly retracting ATPase point mutant PilT^H222A^ on agarose. (A) - (D) Data point is the median of at least three biological replicates and 8 technical replicates each. Error bars are the inter quantile range (IQR) of the distribution.

### The TFP extension and retraction velocities tune the response to substrate stiffness

Our experiments and modelling suggest that PaQa stiffness sensing is determined by the competition between PilA diffusion and the substrate dependent outflux/influx of PilA during TFP extension-retraction. To further test this hypothesis, we investigated the effect of changes in the extension and retraction velocity on stiffness sensing. Both parameters are experimentally accessible by mutations in the ATPase activity of the PilB and PilT motors. Experimentally changing the diffusion of PilA in the inner membrane is not currently experimentally tractable. We first noted that TFP extension velocity on PAA gels, 240 nm/s, is slightly slower than on agarose gels, 350 nm/s, potentially due to a change in the chemical environment. Including this change in the model simulation and changing the adhesion spot size from 1.0 nm on agarose to 0.8 nm on PAA yields the best fit to the experimental P_PaQa_ response (Fig. 5B). These spot sizes are similar to the size, 0.84 nm, that we determined from the TFM and optical trapping experiments on PAA. We also note that the response on PAA is shifted to the right compared to agarose (Fig. 1B), consistent with a slightly smaller spot size of adhesion compared to agarose.

We used two genetic mutants that separately affect the extension and retraction velocity of TFP: PilB^D355N^ reduces the extension velocity on agarose from 350 nm/s to 220 nm/s, and PilT^H222A^ reduces the retraction velocity from 600 nm/s to 250 nm/s. Changing these parameters in the model accurately predicts the experimental P_PaQa_ results, with the exception for very high stiffness with the PilB mutant (Fig. 5). Here, the model slightly overestimates the experimental response similar to PAA. Together, these results demonstrate that the competition of slow PilA diffusion to fast PilA influx during TFP retraction may determine stiffness sensing and the peaked stiffness response.

## Discussion

Here, we experimentally demonstrate that the bacterium *P. aeruginosa* actively measures substrate stiffness using type IV pili and uses this information to express virulence genes in a preferred stiffness range. Our experimental results suggest that the information about substrate stiffness is encoded in temporal changes of the local PilA concentration in the inner membrane during TFP retraction. This conclusion is supported by observations that the concentration of PilA at the base of TFP is dynamic, the concentration dynamics change in a substrate dependent manner, and that genetic mutations that affect the outflux or influx of PilA to/from the membrane affect stiffness sensing. We developed a quantitative mechanical model that consistently explains all these experimental observations.

The dynamic range of stiffness that *P. aeruginosa* can sense covers the entire range of infection sites in the human body [21]. Transcriptional activity of the virulence factor regulator Vfr, which controls hundreds of virulence related genes, is preferentially upregulated at an intermediate stiffness range of 10-100 kPa (Fog. 1B) [47]. Human tissues with stiffnesses in this range include lung, spleen, thyroid, muscle, and skin [48]. This connection between pathogenesis and the mechanical properties of potential infection sites make our discovery of bacterial stiffness sensing interesting for clinical application. In. *P. aeruginosa*, transduction of the force-deformation relationship to transcriptional activity of Vfr relies on the two-component chemosensory system Pil-Chp. Interestingly, many clinically important pathogens like *Pseudomonas*, *Acinetobacter*, and *Stenotrophomonas* encode TFP and Pil-Chp in their genome, making this pathway and the ability to mechanically differentiate different infections sites possibly a widespread and important mechanism during the progression of bacterial infections [49].

Our results suggest that stiffness is sensed by a sensory system that is sensitive to rapid temporal changes in the dynamics of PilA concentration. The chemosensory protein PilJ of the Pil-Chp system is a prime candidate for this purpose. PilJ interacts with the major pilin PilA and PilJ is essential for PaQa activation [25]. Further, chemotaxis systems have evolved to sense temporal changes of ligand concentration, typically with speeds up to 1 s [50]. This timeframe is consistent with the dynamics of the PilA concentration changes we see experimentally and predict in our model. Interestingly, our model predicts a wide distribution of the maximum change in the PilA concentration upon retraction of individual TFP at a given stiffness (Supplementary Fig. 5A). This is a result of the stochastic nature of PilA diffusion and is consistent with the wide distribution of individual P_PaQa_ measurements for a given gel stiffness (Supplementary Fig. 5B). Changes in PilA levels have also been shown to stimulate the activity of the sensor kinase of the PleCD two component system in *C. cresentus* enabling TFP based surface contact sensing [14, 51, 52]. While *C. cresentus* seems to detect changes in the steady state concentration of PilA, our results add an additional layer of complexity where rapid changes in those levels are sensed.

Mechanotransduction allows eukaryotic cells to differentiate substrate stiffness and this process presents a fundamental feature of embryo development and stem cell differentiation into mature tissue [16, 17, 53]. Our experimental results and mechanical model highlight interesting parallels between stiffness sensing in eukaryotes and prokaryotes. Stiffness sensing by the integrin complex for example relies on force dependent stretch and conformational changes that are translated into kinase activity by direct conformation dependent ligand binding. Similarly, we show how stretch induced conformational changes in the TFP might yield a peaked stiffness sensitive activation of the Pil-Chp pathway and its kinase ChpA that stimulates production of the second messenger cAMP. In contrast to the direct activation in eukaryotes, we hypothesize that stiffness sensing in *P. aeruginosa* is converted indirectly from stiffness to ligand concentration that is sensed by a chemotaxis system. Chemotaxis systems are widespread in bacteria and *Pseudomonas* alone has 26 different chemoreceptors [54]. These systems have evolved to sense temporal changes of ligand concentration as the small size of bacteria makes spatial sensing of ligand concentrations challenging. In contrast, the much large eukaryotic cells almost exclusively rely on spatial sensing strategies. Stiffness sensing in eukaryotes is typically reinforced by molecular feedback that recruits proteins to existing focal adhesions and integrin complexes. Similarly, cAMP dependent activation of the transcription factor Vfr yields upregulation of TFP genes yielding a feedback of the stiffness sensing components.

Together, our results demonstrate that stiffness sensing is a conserved ability in pro- and eukaryotes. Our model in conjunction with genetic tests suggest that the stiffness sensing mechanism is similar between both trees of life and that their specific differences have evolved reflecting the different molecular prerequisites, i.e., spatial or temporal gradient sensing. TFP are broadly conserved, and thus bacterial stiffness sensing is potentially a physiologically important feature.

## Materials and Methods

### Strains and Growth conditions

Information on cloning, plasmids, and primers used in this study can be found in the SI Appendix, Materials and Methods and Tables S2–S4.

*P. aeruginosa* PAO1 was grown in liquid lysogeny broth (LB) Miller (Difco) in a floor shaker, on LB Miller agar (1.5% Bacto Agar), on Vogel-Bonner minimal medium (VBMM) agar (200 mg/l MgSO4 7H2O, 2 g/l citric acid, 10 g/l K2HPO4, 3.5 g/l NaNH4HPO4 4 H2O, and 1.5% agar), and on no-salt LB (NSLB) agar (10 g/l tryptone, 5 g/l yeast extract, and 1.5% agar) at 30 °C (for cloning) or at 37 °C. *Escherichia coli* S17 was grown in liquid LB Miller (Difco) in a floor shaker and on LB Miller agar (1.5% Bacto Agar) at 30 °C (for cloning, see below) or at 37 °C. Antibiotics were used at the following concentrations: 200 μg/mL carbenicillin in liquid (300 μg/mL on plates) or 10 μg/mL gentamycin in liquid (30 μg/mL on plates) or 10 μg/mL anhydrotetracycline in liquid for *P. aeruginosa*, and 100 μg/mL carbenicillin in liquid (100 μg/mL on plates) or 30 μg/mL gentamycin in liquid (30 μg/mL on plates) for *E. coli*.

## Acknowledgements

The authors like to thank Ned Wingreen, Chenyi Fei, Joseph Sanfilippo, Ben Bratton, Nick Martin, Courtney Ellison, Geoff Vrla, and Benedikt Sabass for stimulating discussions and advice. We further like to thank Gary Laevski from the Princeton Molecular Biology Confocal Microscopy Core Facility and Nikon Center of Excellence.

## Author Contributions

All authors designed the research. M.D.K. performed all experiments, analyzed the data, developed the model, and performed simulations. E.H. modelled substrate deformations and analyzed TFM data. M.D.K., J.W.S. and Z.G. wrote the paper.

## Supplementary Appendix

### Supplementary Methods

#### Strain construction

The *pilB*^D355N^ and *pilT*^H222A^ chromosomal point mutations were generated using two-step allelic exchange [55]. Briefly, the cloning vectors were created by digesting the pEXG2 backbone with the HindIII HF restriction enzyme (NEB). The inserts were created the following: for *pilB*^D355N^, the 500 bp flaking regions of the mutation site were PCR amplified using primers pilB_D355N P1/2 and P3/4. Primers pilB_D355N_P2/3 are reverse complements to each other and contain the point mutation. Both fragments were then inserted into the backbone by Gibson assembly according to the manufacturers protocol using the NEB Gibson Assembly Master Mix (NEB). For *pilT*^H222A^, an insert was created to complement back into a Δ*pilTU* mutant. This strategy makes screening for the correct point mutation on the chromosome faster compared to the direct mutation strategy used for *pilB*^D355N^. The regions starting 1000 bp upstream of *pilT* to the point mutation site and from the point mutation site to 1000 bp downstream of *pilU* were PCR amplified using primers pilT_H222A_P1/2 and pilT_H222A_P3/4. Primers pilT_H222A2/3 are reverse complements to each other and contain the point mutation. Both fragments were joined using sewing PCR using the flanking primers, subsequently digested using HindIII HF, and ligated into the pEXG2 backbone. Both cloning vectors were electroporated into *E. coli* and the correct mutation was PCR screened and sanger sequenced using primers pEXG2_Ver1/2. For mating, 1.5 ml *E. coli* containing the vector were grown to OD 0.5. The *P. aeruginosa* parental strain was grown overnight, and 0.5 ml culture was diluted 1:2 into fresh LB and incubated for 3 hours at 42 °C. Both cultures were concentrated into 100 μl and spotted onto an LB agar plate and incubated overnight at 30 °C. The puddle was scrapped off, resuspended into 150 μl PBS, spread onto a VBMM plate containing 30 μg/ml gentamycin and incubated 24 hours at 37 °C. Six single colonies from the VBMM plate were struck onto NSLB and incubated for 24 hours at 30 °C. Several single colonies from the NSLB plate were screened for the correct mutation using PCR amplification with the flaking primers and confirmed using sanger sequencing.

#### Preparation of hydrogels

<u>Agarose gels</u> were prepared according by microwaving and subsequent cool down to 60 C in a water bath. Agarose was prepared freshly every day. An approximately 3 mm thick gel was made with the help of adhesive silicone isolators (Grace Bio-Labs) between a microscope slide and a coverslip, cured for 20 min and dried for 3 min after removal of the cover slip to remove excess liquid on the surface and avoid planktonic cells, i.e., to increase interaction with the surface.

<u>Polyacrylamide (PAA) gels</u> were either prepared from commercially available acrylamide (AA) and bisacrylamide (BIS) solutions (Biorad) (for gels with G’ < 130 kPa) or solid AA and BIS by dissolving in ddH2O (for gels with G’ > 130 kPa) according to the ratios for total AA + BIS content 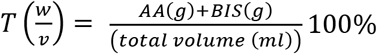 and crosslinker content 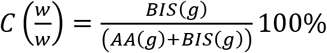 given by Supplementary Table 1 [28]. For each gel, 50ml of total solution was prepared and polymerized adding 250 μl 10% APS (Biorad) and 75 μl Temed (Biorad). After mixing AA and BIS, the solution was vacuum degassed for 10 min to remove oxygen which hampers polymerization and polymerized for 60 min at room temperature in a 150 mm petri dish with another 145 mm petri dish on top to get a planar surface of an approximately 5 mm thick gel. Crosslinking of PAA is an exothermal process which becomes apparent for gels with T > 20%. Such gels were polymerized in a water bath with the addition of small amounts of ice to keep the entire reaction at room temperature. Gels were then washed twice with LB medium and soaked overnight in LB. One centimeter of the edge of each gel was cut off before washing because gels swell slightly during soaking with growth medium. The next day, liquid was removed and the gel surface was dried for 20 min in a sterile flow hood. This removes excess liquid on the surface and avoids planktonic cells, i.e., increases interaction with the surface. Gels were prepared freshly every day.

**Supplementary Table 1.** Polyacrylamide gel preparation. Stiffnesses estimated from [28].

|  |  |  |  |  |  |  |  |  |  |  |  |  |
| --- | --- | --- | --- | --- | --- | --- | --- | --- | --- | --- | --- | --- |
| Stiffness<br>G' (kPa) | 0.21±0.04 | 0.73±0.18 | 1.7±1.1 | 4.7±0.6 | 16±1 | 45±5 | 90±30 | 132±23 | 222±50 | 311±50 | 410±50 | 600±100 |
| % T | 3.1 | 5 | 7.6 | 7.7 | 12.1 | 12.6 | 16.4 | 20 | 25 | 29 | 33 | 43 |
| % C | 2 | 1 | 0.7 | 2.6 | 1.2 | 4.8 | 4.9 | 5 | 6 | 7.5 | 9 | 7 |

#### Microscopy, sample preparation, and data analysis

##### TFP imaging

A commercial Nikon Ti-E TIRF microscope was used in HILO mode where the TIRF angel is set to slightly below the critical angle. This configuration reduces background and allows to image TFP dynamics longer and with higher contrast compared to regular EPI fluorescence imaging. The microscope was used with perfect focus, a 100x NA 1.49 Apo TIRF lens (Nikon), an EMCCD camera (iXon Ultra DU-897U, Andor), a stage top incubator (INU, Tokai Hit) set to 37° C and controlled by Nikon Elements software. TFP were fluorescently labeled as described previously [34]. TFP dynamics were analyzed manually in ImageJ.

##### P_PaQa_ timelapse imaging

Movies were taken on a Nikon Ti-E inverted microscopes controlled by Nikon NIS elements software using a 40x NA 0.95 Plan Apo Phase lens, an Andor Clara CCD or Hamamatsu Orca Flash camera, equipped with a environmental chamber (In Vivo Scientific) for temperature control set to 37° C. For all experiments, a stationary phase overnight culture (> 18 hrs) in LB was diluted 1:2000 in fresh LB and grown for 120 min at 37 C in a shaking incubator. 1.5 μl of cell suspension was then spread on an approx. 10 x 10 mm large gel pad. The cell suspension was given 3-5 min to allow drying. Excess liquid on the gel creates a thin liquid film between gel and cover slip and a significant number of cells swims rather than adhering to the surface. The pad was then transferred to a glass bottom dish (Matek) and 200μl of water was distributed to 8-10 small droplets at the edge of the dish to keep the sample moisturized and to minimize sample degradation over time. Each gel was imaged every 20 min for 4 hours. Each experiment is a multi-point acquisition of typically 8 different fields of view (FOV) on the same gel, each FOV containing 10 – 40 cells at the beginning. At least three independent replicates per substrate / stiffness were acquired. Individual cells were segmented using a custom written, threshold-based segmentation algorithm based on the constitutive reporter (P_rpoD_::mkate2) image to create a binary mask. This allowed estimating the mean background and fluorescence signal of both channels, which were analyzed as the ratio mVenus2 / mKate2. The ratio was analyzed for each time point and the median and inter quantile ranges (IQR) for the distribution at each time point was calculated.

##### Optical trapping

The custom built optical trapping setup consists of a 5 W 1064 nm laser (Spectra Physics) focused in the focal plane of a 100x NA 1.49 Apo TIRF lens (Nikon) to create an optical trapping potential [56]. For detection, we used back focal plane interferometry using a 850nm laser (QPhotonics), a position sensitive photodetector (New Focus), and an analog to digital conversion card for data acquisition (National Instruments). The trap and detection focus were positioned onto each other using a tip-tilt piezo mirror (Mad City Labs). Samples were positioned using a three-axis piezo stage (Mad City Labs). Brightfield images were acquired using a water cooled EMCCD camera (iXon Ultra DU-897, Andor). The entire microscope was controlled using custom written software in National Instruments LabView. Experiments were performed using a custom built, laser-cut incubation chamber and a PID temperature controller (In Vivo Scientific) set to 37° C. For experiments, an overnight culture was diluted 1:50 in fresh LB and grown for 3 hrs at room temperature. 20 ul of cell suspension was mixed 1:10000 with 532 nm polystyrene beads (Bangs Labs) and flushed by capillary action into a tunnel slide made by taping a coverslip to a microscope slide using double sided sticky tape. A trapped bead was then brought close to the pole of individual cells and we waited until the bead was displaced by TFP retraction towards the cell. Individual traces of bead displacements were analyzed in Igor Pro (Wavemetrics). Traces were calibrated using the thermal fluctuations method [56]. As shown in Supplementary Fig. 3B, an exponential fit to the bead position and force was used to estimate the force free velocity (Fig. 3D) and stalling force (Fig. 3C). To obtain the force-velocity relationship of TFP retraction, the high frequency bead displacements and forces were down-sampled/smoothed and differentiated.

##### Traction force measurements (TFM)

A commercial Nikon Ti-E TIRF microscope was used with a 100x NA 1.49 Apo TIRF lens (Nikon), an EMCCD camera (iXon Ultra DU-897U, Andor), a stage top incubator (INU, Tokai Hit) set to 37° C and controlled by Nikon Elements software. PAA gels were made as described above with the addition of 2.5 μl of red-orange (565/580) and dark red (660/680) 0.04 μm carboxylated beads (Thermo Fisher FluoSpheres) per 500 μl gel solution. Hydrogels were polymerized between two chemically modified coverslips [57]. A 22 x 40 mm coverslip (Fisherbrand) was methacrylate silanized by submerging plasma cleaned coverslips in 2% 3-(trimethoxysilyl) propylmethacrylate in 95% ethanol for 10 minutes. Hydrophobic round 12 mm coverslip (Fisherbrand) were prepared by submerging coverslips in Sigmacote (Sigma) for 10 minutes. All coverslips were rinsed three times in 100% ethanol and dried upright on kim wipes. About 10 ul of unpolymerized gel was added between both coverslips resulting in an approximately 50 um tall gel. Gels were polymerized for 30 minutes at room temperature, followed by removal of the hydrophobic coverslip. Gels were washed three times and then incubated for 30 minutes in EZ rich medium. For experiments, overnight cultures were diluted 1:200 and grown to mid log phase at 37° C shaking in LB. Excess liquid was removed carefully from the gel using kim wipes and 1.5 μl of cell suspension was added. Cells were incubated on the gel for 5 minutes before excess liquid was removed carefully using kim wipes. A top 22 x 22 mm coverslip was added and sealed with Valap. Bead displacement caused by TFP retraction were analyzed manually in ImageJ.

##### Fluorescence recovery after photobleaching (FRAP)

We used a custom built FRAP microscope. To allow quasi simultaneous widefield EPI fluorescence imaging and localized photo bleaching, we split a 488 nm laser (Coherent) into two separate beam paths using a motorized flip mirror (Thorlabs). One beam path was focused in the back focal plane of a 100x NA 1.49 Apo TIRF lens (Nikon) to allow even illumination of the focal plane. A second beam path was focused in the focal plane of the objective lens to enable diffraction limited localized photo bleaching. The flip mirror allows the sequential use of both path and flipping within less than 1 s. Samples were positioned using a three axis piezo stage (Mad City Labs) and images wee acquired using a water cooled EMCCD camera (iXon Ultra DU-897, Andor). The entire microscope was controlled using custom written software in National Instruments LabView. Experiments were performed using a custom built, laser-cut incubation chamber and a PID temperature controller (In Vivo Scientific) set to 37° C. For experiments, the *pilA*^A86C^ strain was grown and fluorescently labeled as described above. Cells were imaged fro a few seconds to establish a pre-bleach fluorescence baseline, then bleached for 100 ms, and immediately imaged again at 1 s frame rate. To obtain the fraction of immobile fluorescence molecules, the experimental data have to be corrected for both bleaching due to FRAP and bleaching during fluorescence imaging. The loss of fluorescence intensity due to FRAP bleaching was corrected by analyzing the total fluorescence of the entire cell immediately before and after the FRAP bleaching event, and images taken after FRAP bleaching were corrected respectively. Photobleaching due to imaging was corrected by analyzing the total fluorescence intensity of cells outside of the FRAP bleaching spot, fitted by an exponential which was applied to all images. The rise of fluorescence at the FRAP bleaching spot was the nanalyzed over time and fitted with an exponential. This yields the recovery plateau (immobile fraction) for t →∞ and the time constant *τ* of the recovery. The diffusion coefficient *D* can then be calculated by 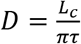 using the length of the cell *L_c_* [35].

#### Estimation of hydrogel stiffness and pore size distribution

Data on agarose and PAA stiffness *G*’ and pore size *D* of the gel network are reported by different groups but vary depending on the experimental method used. For PAA stiffness as a function of T and C, we used the data provided in the Supplementary Information of the meta study [28] and averaged stiffness values that are +/- 10 % of the same composition (typically 2 - 4 reported values). Chui et al. [30] report a PAA pore size < 10 nm even for the lowest concentration gels using NMR while Holmes and Stellwagen [31] report a pore size of up to 150 nm for soft gels and < 20 nm for stiff gels using electrophoresis. We tried to validate one or the other by embedding uncoated beads of varying size in PAA hydrogels and found that 40 nm beads are still somewhat diffusive in very soft gels (230 Pa) but not anymore in slightly stiffer gels of 1 kPa. Similarly, 60 nm beads are completely immobilized in the network of a 230 Pa gel. Hence, we conclude that PAA gel pore size is smaller than 60 nm in all cases. To get a more realistic estimate, we combined the data by [30, 31] for high stiffness with our estimate for soft gels and used a power law fit to get a functional relation between stiffness and pore size: *D* ≈ 9.5 + 26 *G* - 0.43, where *G* is in kPa in *D* in nm.

For agarose, we mainly used the power law estimate *G* ~ *c*^1.8^ by [29] that links the shear modulus with the w/w concentration *c* of agarose. This estimate fits well two other publications [58, 59]. Data on agarose pore size varies among different reports and we decided to take all the available data [30, 32, 33] and empirically found that a double exponential fit function to all of the available data describes the pore size distribution as a function of agarose concertation *c* well. Consistent with the estimate that 0.2 % agarose gels have a pore size > 1 μm, we observed few cases where single *Pseudomonas* cells were able to move through 0.20 % agarose gels, although very slowly and in a non-persistent way.

#### Estimation of the total number of PilA proteins in a single cell

From the fluorescent PilA concentration dynamics experiment described in the main text, we measured a total decrease of PilA fluorescence in the entire cell of 10% for the pilus that extend for 5 μm. The packing of PilA in the TFP fiber of the closely related *Pseudomonas K* strain (PAK) is approximately 4 PilA monomers per 4 nm of TFP fiber [60]. This equals to 20000 PilA monomers for a 5 μm long TFP or 200000 PilA monomers total per cell. Based on this analysis we can model the local loss of PilA at the TFP base during extension and the local source of PilA at the base during retraction.

#### Surface deformation, TFP retraction, and PilA diffusion models

The surface displacement *Δx* generated by a force *F* applied uniformly in a circular area with radius *R* is given by

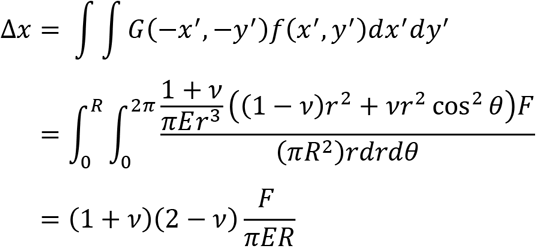

where *v* = 0.5 is the Poisson’s ratio of PAA and *E* = 3*G* is the Young’s modulus (see Supplementary Fig. 3A). This yields the force-deformation relation 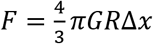. Based on our optical trapping experiments, the force-velocity relation of TFP retraction can be described by *ν*(*F*) = *ν*_0_ – *ν_f_* · *F*. The substrate deformation is coupled to TFP retraction by 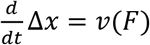. For a polymer with *N* = *N*_0_(*t*) + *N*_*s*(_*t*) monomers, the transition rates of folded *N*_0_(*t*) to unfolded *N_s_*(*t*) protein can be described by 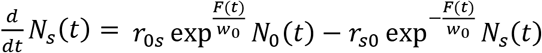 and 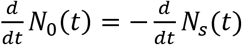, where *r*_0s_ and *r*_s0_ are the force free equilibrium transition rates from unstretched to stretched, and stretched to unstretched, respectively, and *w*_0_ = *k_B_T* / *δ_a_* reflects the shape of the potential along the reaction coordinate *δ_a_* [61]. The current rest length of the TFP fiber is given by *l*_*p*0_ – Δ*x*(*t*), where *l*_*p*0_ is the maximum TFP length immediately prior to the start of retraction. At the same time, stretching of PilA monomers yields the length of the TFP fiber *l_p_*(*t*) = *a*_0_*N*_0_(*t*) + *a_s_N_s_*(*t*), where *a*_0_ = 1 nm and *a_s_* = 3*a*_0_ are the length of a unstretched and stretched PilA monomer in the fiber, respectively [38, 60]. Since 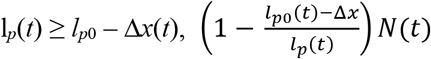 monomers can be removed from the TFP fiber and retracted into the cell, given the maximum catalytic ratio (600nm/s ≡ 600 PilA/s) determined by ATP hydrolysis is not violated.

We performed numerical simulations with 1 ms time steps of this coupling of substrate deformation, TFP retraction, and PilA unfolding to determine the influx of PilA as a function of time and substrate stiffness.

The outflux of PilA monomers during TFP extension is trivial and given by *v_ext_* · 1 PilA/nm = 350 PilA / nm for WT with the TFP extension velocity *v_ext_* = 350 nm/s. This sequential outflux and influx of PilA results in a PilA loss function which we coupled to a Brownian dynamics simulation of PilA diffusion in the inner membrane. Here, we modeled the inner membrane as a one-dimensional space with circular boundary conditions equivalent to the cross-sectional dimension of an average cylindrical cell with spherical endcaps with a length of 3 μm and a diameter of 0.5 μm. The diffusion of each individual monomer can then be described numerically by 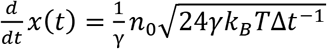, where *n*_0_ is a random number drawn from a uniform distribution in the interval −0.5 ≤ *n*_0_ ≤ 0.5, *γ* = *k_B_T* / *D*_PilA_ is the friction of PilA in the inner membrane given by the Einstein relation and the experimentally obtained diffusion coefficient *D* = 0.22 μm^2^/s, and Δ*t* = 1 ms is the time step of the numerical simulation [62].

We determined most of the parameters of this model experimentally and inferred *a*_0_ and *a_s_* from published data (see above). Only the variables *r*_0s_, *r*_s0_, *w*_0_, and *R* remain as free fit parameters. These parameters were varied iteratively until we obtained a best match with the experimental data for WT cells on agarose. The best fit values are: *r*_0s_ = 1 · 10^-6^ s^-1^, *r*_s0_ = 1000 s^-1^, *w*_0_ = *k_B_T*/ *δ_a_* = 3.4 pN, equivalent to *δ_a_* =1.22 nm, and *R* = 1 nm.

**Supplementary Table 2.** Strains used in this study.[63, 64]

| Strain | Description | Reference |
| --- | --- | --- |
| *E. Coli*<br>S17 | Wild-type, used for cloning and conjugation |  |
| *P. aeruginosa*<br>PAO1 | Wild-type from Manoil transposon mutant library |  |
| ZG 1332 | Wild-type PAO1 with pPaQa-mVenus plasmid | Ref 25. |
| ZG 1586 | PAO1 *pilA* -A86C chromosomal point mutation | Ref 34. |
|  | PAO1 *pilB* -D355N chromosomal point mutation. | This study |
|  | PAO1 *pilB* -D355N *pilA* -A86C chromosomal point mutations | This study |
|  | PAO1 *pilB* -D355N chromosomal point mutation with pPaQa-mVenus plasmid | This study |
|  | PAO1 *pilT* -H222A chromosomal point mutation | This study |
|  | PAO1 *pilT* -H222A *pilA* -A86C chromosomal point mutations. | This study |
|  | PAO1 *pilT* -H222A chromosomal point mutation with pPaQa-mVenus plasmid | This study |
| ZG 1183 | PAO1 $\Delta pilTU$, in frame deletion of pilTU | Ref 62. |
| | PAO1 $\Delta pilTU$ with pPaQa-mVenus plasmid | Ref 25. |
| ZG 1186 | PAO1 $\Delta vfr$, in frame deletion of vfr | Ref 63. |
| | PAO1 $\Delta vfr$ with pPaQa-mVenus plasmid | This study |

**Supplementary Table 3.** Plasmids used in this study.

| Plasmid | Description | Reference |
| --- | --- | --- |
| pEXG2 | Vector for generating deletion mutants and point mutants | Ref 54. |
| ZG 1658 - pEXG2-*pilA* -A86C | Introduces A86C point mutation in PAO1 *pilA* | Ref 34. |
| ZG XXXX - pEXG2-*pilB* -D355N | Introduces D355N point mutation in PAO1 *pilB* | This study. |
| ZG XXXX - pEXG2-*pilT* -H222A | Introduces H222A point mutation in PAO1 *pilT* | This study. |
| ZG 1190 - pPaQa-mVenus | pUCP18 backbone with PPaQa::mVenus2, PrpoD::mKate2 | Ref 25. |

**Supplementary Table 4.**
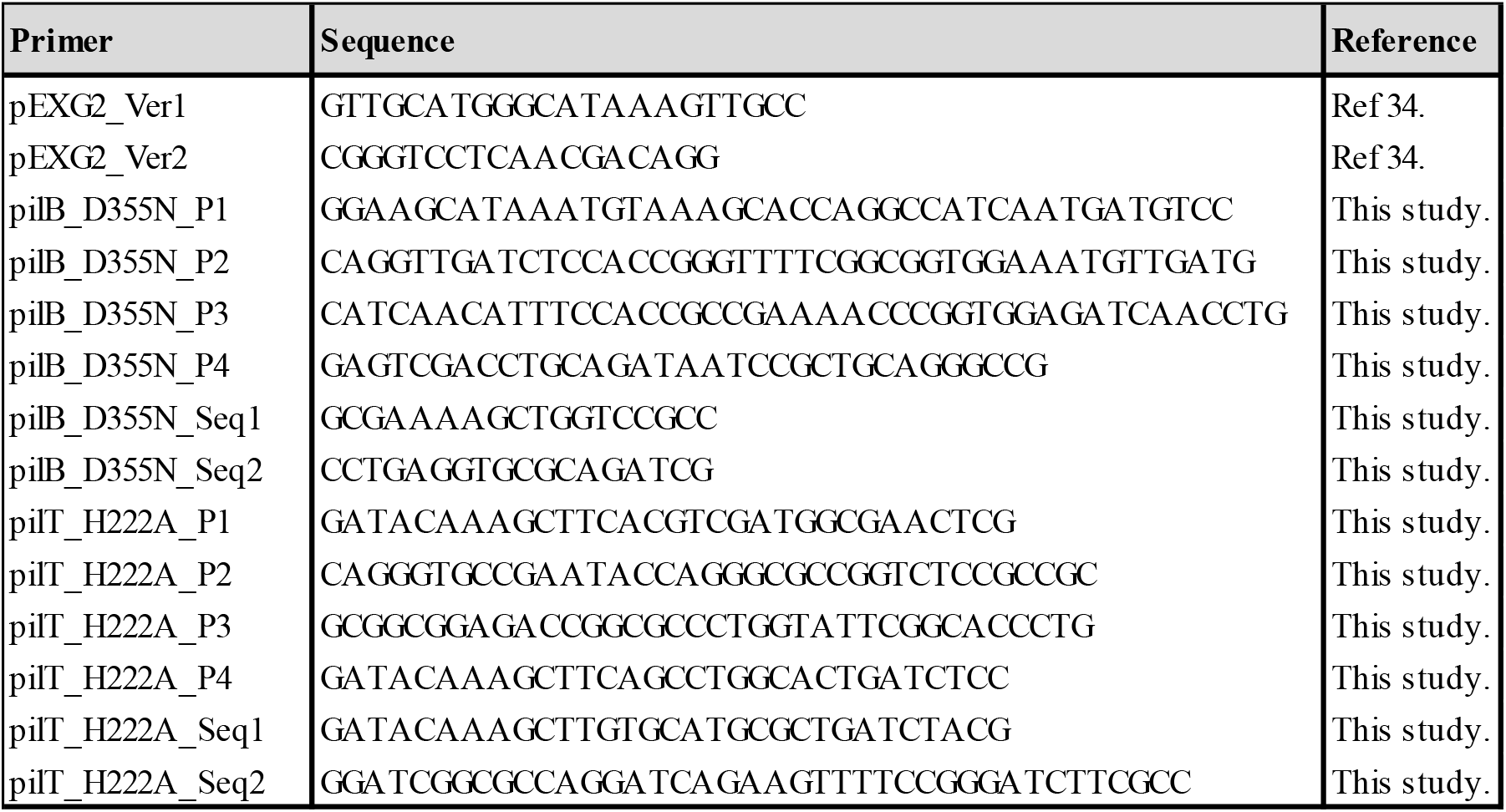
Primers used in this study.

### Supplementary Figures

**Supplementary Fig. 1.**
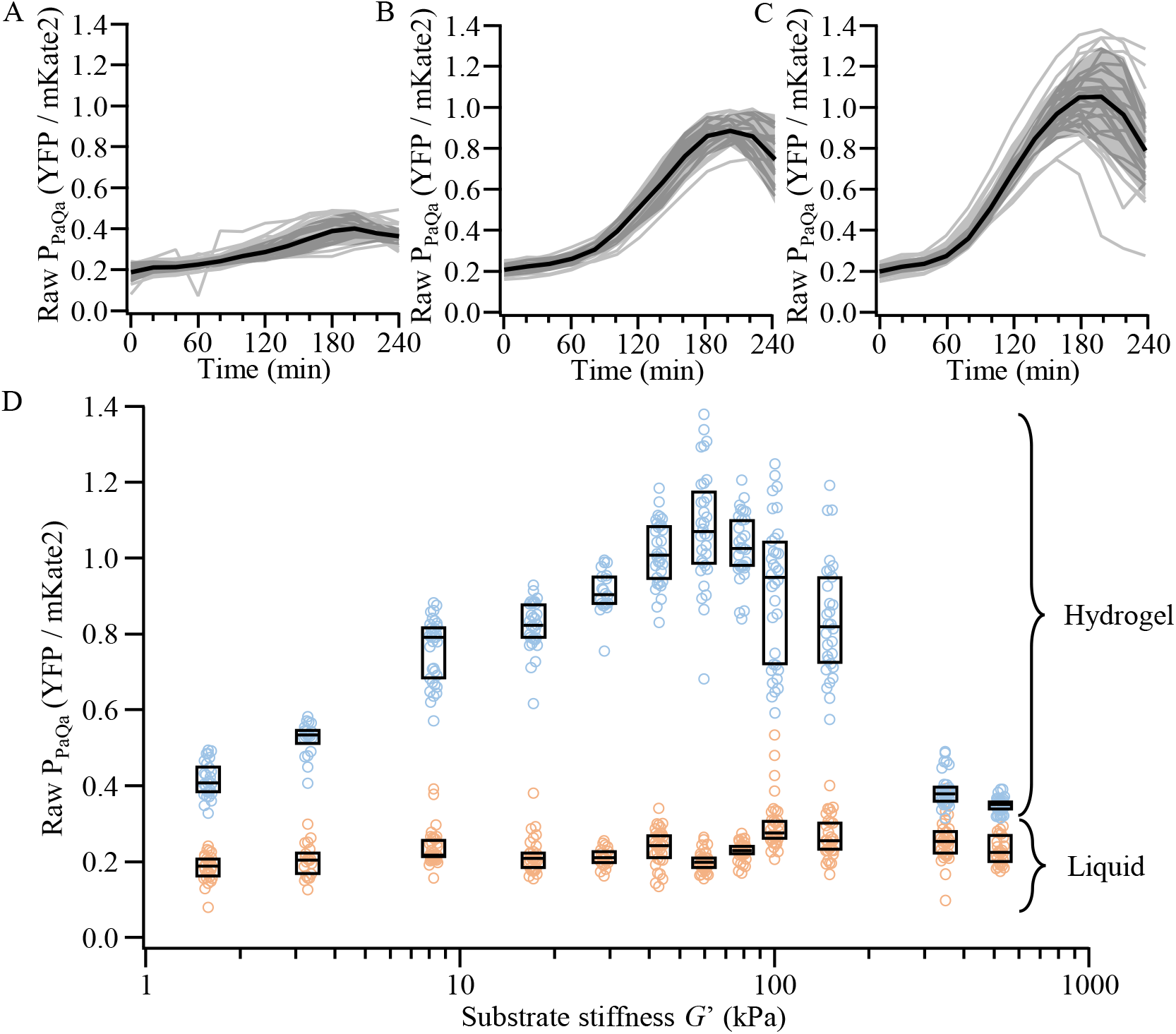
Experimental response of P_PaQa_ to the stiffness of different agarose hydrogels. (A) The responses of 27 technical replicates from 3 biological replicates on 0.2 % agarose (≈ 1.5 kPa). (B) The responses of 19 technical replicates from 3 biological replicates on 1.0 % agarose (≈ 30 kPa). (C) The responses of 31 technical replicates from 3 biological replicates on 1.5 % agarose (≈ 60 kPa). (A) – (C) The median and IQR are shown as solid black line and gray shaded area. (D) The distributions of the raw P_PaQa_::YFP / P_rpoD_::mKate2 fluorescence intensity on various agarose hydrogels. “Liquid” is the zero time point and “Hydrogel” is the maximum response at 180 min after the experiment started. Boxes represent the median and 25/75% quantiles.

**Supplementary Fig. 2.**
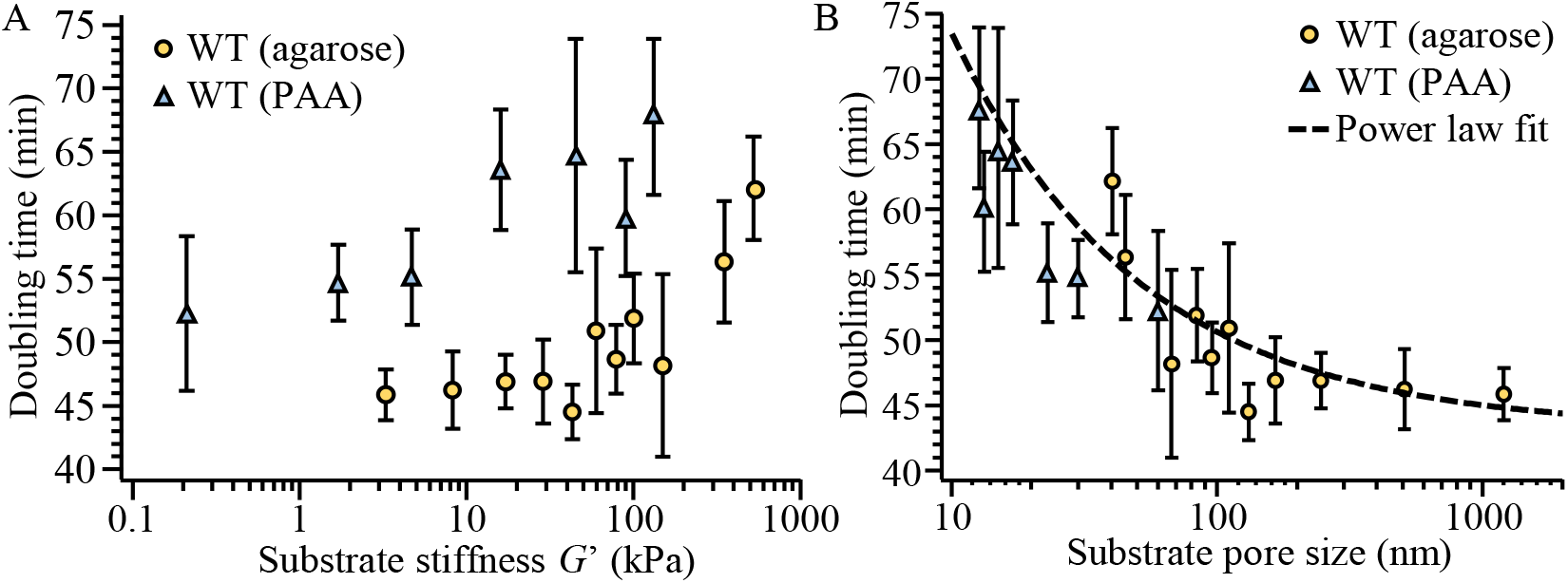
Experimentally obtained doubling time of cells grown on different agarose and PAA hydrogels. (A) The doubling time as a function of substrate stiffness does not collapse onto each other for both gel types, indicating that the doubling time is not a function of substrate stiffness. (B) The doubling time as a function of substrate pore size does collapse onto each other for both gel types, indicating that the doubling time is a function of substrate stiffness.

**Supplementary Fig. 3.**
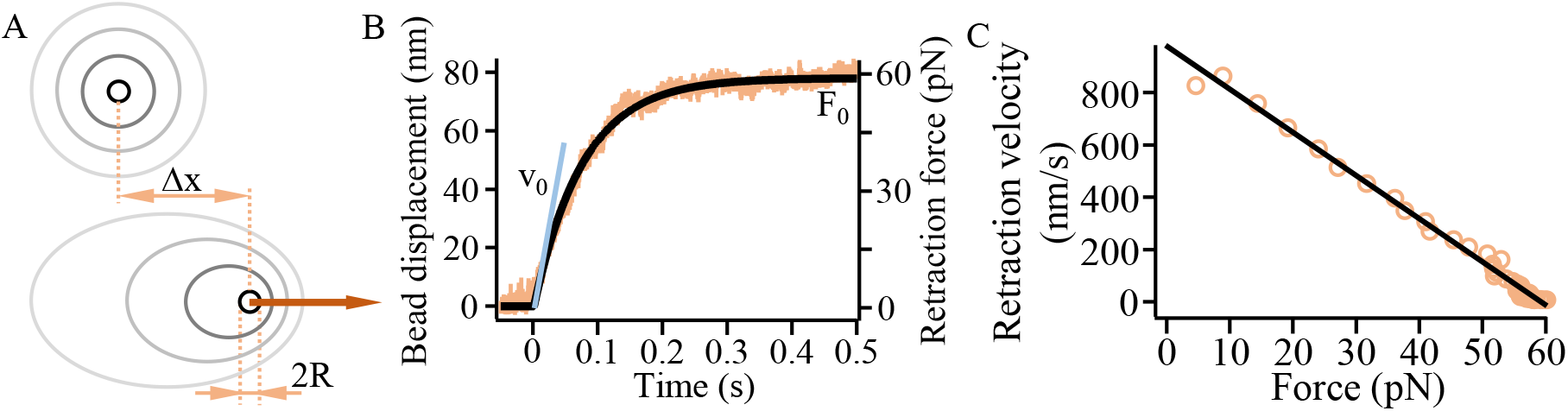
Surface deformation and optical trapping. (A) Sketch of the surface deformation of a substrate. The force *F* is uniformly applied in the circular area with diameter 2*R*, yielding the deformation Δ*x* along *F*. (B) Exemplary displacement of an optical trapped bead by the retraction of a TFP. (C) Relation of the retraction velocity as a function of the retraction force from B).

**Supplementary Fig. 4.**
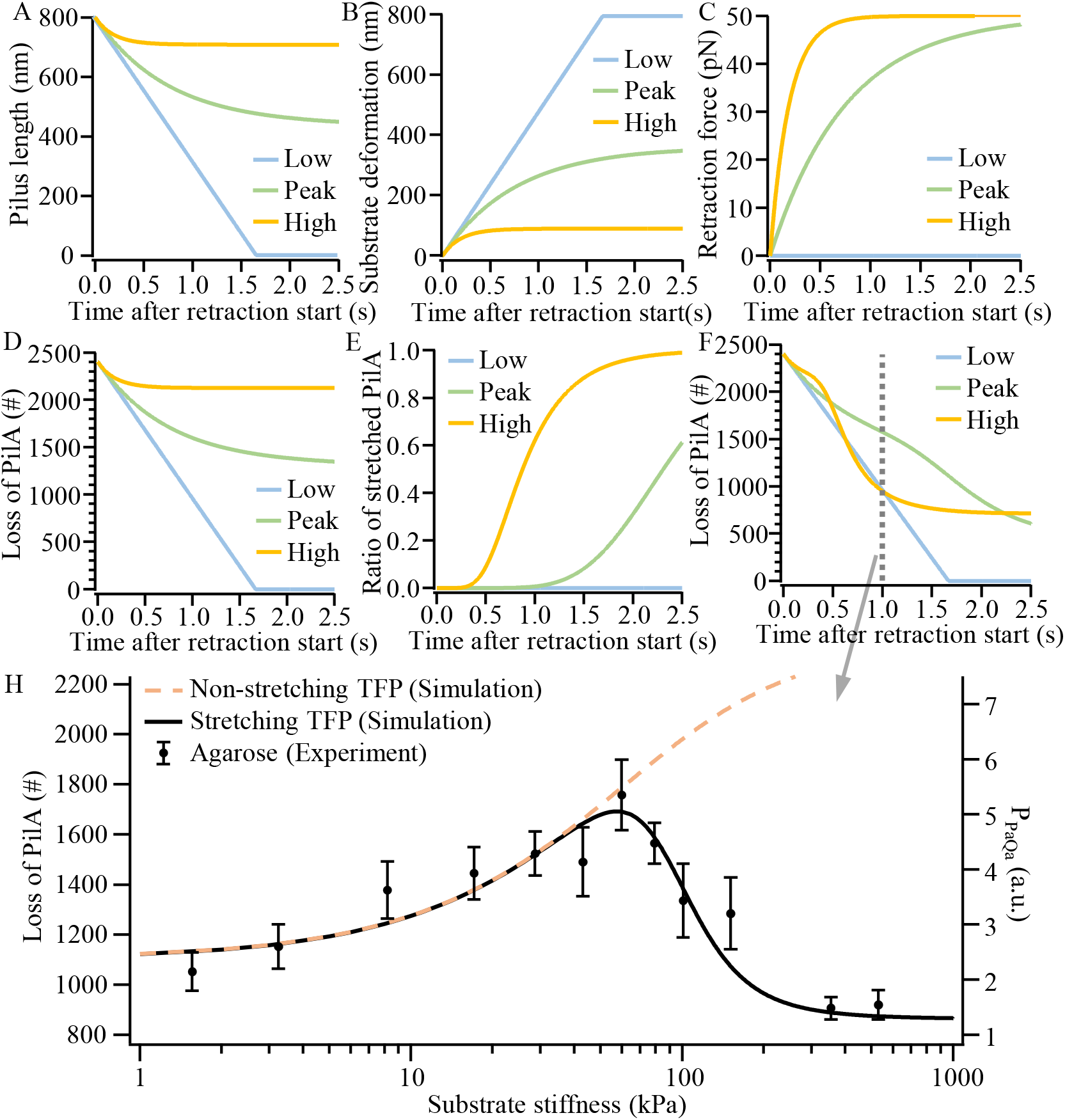
Mechanical surface deformation TFP retraction model. (A) - (F) Model result for a typical 0.8 μm long TFP on three different substrates (Low = 1 kPa, Peak = 60 kPa, High = 200 kPa). (A) Pilus length. (B) Substrate deformation. (C) Retraction force needed to deform the substrate. (D) Loss of PilA in the inner membrane for the non-stretching model. This value starts high because the TFP is fully extended at *t* = 0. (E) The ratio of stretched PilA to all PilA in the TFP fiber in the stretching model. (F) Loss of PilA in the inner membrane for the stretching model. This value starts high because the TFP is fullt extended at *t* = 0. (H) Loss of PilA in the inner membrane at T = 1 s after the start of retraction as a function of substrate stiffness for the non-stretching model (red dashed line), the stretching model (solid black line), and the experimental P_PaQa_ data for WT on agarose.

**Supplementary Fig. 5.**
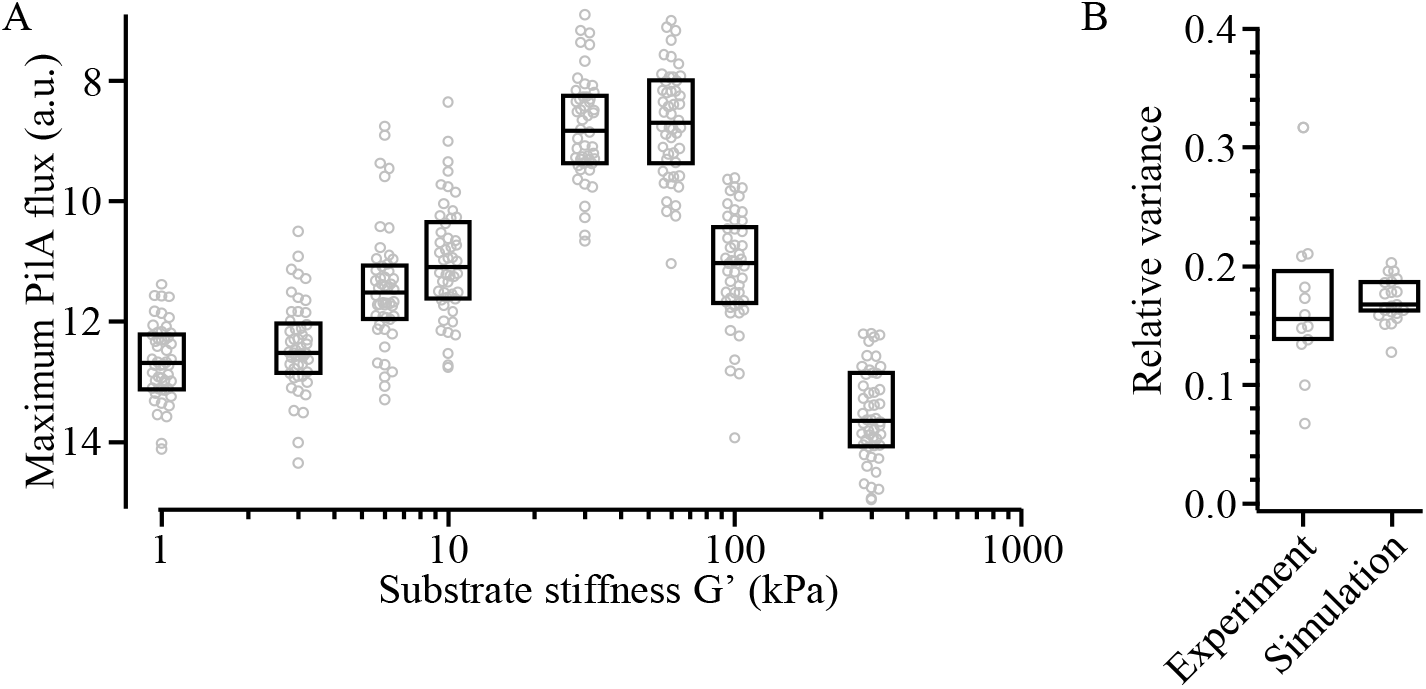
Variance in the response to stiffness for individual events. (A) Variance of individual model simulations with identical parameters. Compare to Supplementary Fig. 1C. (B) Comparison of variance between simulations in A) and experiments in Supplementary Fig. 1C. Each data point represents the relative variance of all simulations/experiments at one stiffness. The relative variance of the experiments of one stiffness is calculated as the standard deviation of all datapoints of that stiffness relative to the difference of the lowest to the largest median of the responses to all stiffness (e.g., for the simulations show in A), 13.5 – 9 = 4.5). Boxes represent the median and 25/75% quantiles.

## References

1. Persat, A., et al., The mechanical world of bacteria. Cell, 2015. 161(5): p. 988–997.

2. Jiang, Z., et al., Searching for the Secret of Stickiness: How Biofilms Adhere to Surfaces. Frontiers in microbiology, 2021. 12(1642).

3. O’Toole, G.A. and G.C. Wong, Sensational biofilms: surface sensing in bacteria. Current opinion in microbiology, 2016. 30: p. 139–146.

4. Chawla, R., et al., A skeptic’s Guide to Bacterial Mechanosensing. Journal of Molecular Biology, 2020. 432(2): p. 523–533.

5. Graham, K.J. and L.L. Burrows, More than a feeling: microscopy approaches to understanding surface-sensing mechanisms. Journal of Bacteriology, 2020. 203(6): p. e00492–20.

6. Wong, G.C., et al., Roadmap on emerging concepts in the physical biology of bacterial biofilms: from surface sensing to community formation. Physical biology, 2021.

7. Laventie, B.-J. and U. Jenal, Surface sensing and adaptation in bacteria. Annual review of microbiology, 2020. 74: p. 735–760.

8. Bruni, G.N., et al., Voltage-gated calcium flux mediates Escherichia coli mechanosensation. Proceedings of the National Academy of Sciences, 2017. 114(35): p. 9445–9450.

9. Webster, S.S., et al., Interaction between the type 4 pili machinery and a diguanylate cyclase fine-tune c-di-GMP levels during early biofilm formation. Proceedings of the National Academy of Sciences, 2021. 118(26).

10. Hug, I., et al., Second messenger–mediated tactile response by a bacterial rotary motor. Science, 2017. 358(6362): p. 531–534.

11. Hershey, D.M., Integrated control of surface adaptation by the bacterial flagellum. Current opinion in microbiology, 2021. 61: p. 1–7.

12. Lele, P.P., B.G. Hosu, and H.C. Berg, Dynamics of mechanosensing in the bacterial flagellar motor. Proceedings of the National Academy of Sciences, 2013. 110(29): p. 11839–11844.

13. Burrows, L.L., Pseudomonas aeruginosa twitching motility: type IV pili in action. Annual review of microbiology, 2012. 66: p. 493–520.

14. Ellison, C.K., et al., Obstruction of pilus retraction stimulates bacterial surface sensing. Science, 2017. 358(6362): p. 535–538.

15. Vogel, V. and M. Sheetz, Local force and geometry sensing regulate cell functions. Nature Reviews Molecular Cell Biology, 2006. 7(4): p. 265–275.

16. Discher, D.E., P. Janmey, and Y.-l. Wang, Tissue cells feel and respond to the stiffness of their substrate. Science, 2005. 310(5751): p. 1139–1143.

17. Engler, A.J., et al., Matrix elasticity directs stem cell lineage specification. Cell, 2006. 126(4): p. 677–689.

18. Jaalouk, D.E. and J. Lammerding, Mechanotransduction gone awry. Nature Reviews Molecular Cell Biology, 2009. 10(1): p. 63–73.

19. DuFort, C.C., M.J. Paszek, and V.M. Weaver, Balancing forces: architectural control of mechanotransduction. Nature Reviews Molecular Cell Biology, 2011. 12(5): p. 308–319.

20. Harper, C.E. and C.J. Hernandez, Cell biomechanics and mechanobiology in bacteria: Challenges and opportunities. APL bioengineering, 2020. 4(2): p. 021501.

21. Bodey, G.P., et al., Infections caused by Pseudomonas aeruginosa. Reviews of infectious diseases, 1983. 5(2): p. 279–313.

22. O’Toole, G.A. and R. Kolter, Flagellar and twitching motility are necessary for Pseudomonas aeruginosa biofilm development. Molecular Microbiology, 1998. 30(2): p. 295–304.

23. Siryaporn, A., et al., Surface attachment induces Pseudomonas aeruginosa virulence. Proceedings of the National Academy of Sciences, 2014. 111(47): p. 16860–16865.

24. Laventie, B.-J., et al., A surface-induced asymmetric program promotes tissue colonization by Pseudomonas aeruginosa. Cell host & microbe, 2019. 25(1): p. 140–152. e6.

25. Persat, A., et al., Type IV pili mechanochemically regulate virulence factors in Pseudomonas aeruginosa. Proceedings of the National Academy of Sciences, 2015. 112(24): p. 7563–7568.

26. Maier, B., et al., Single pilus motor forces exceed 100 pN. Proceedings of the National Academy of Sciences, 2002. 99(25): p. 16012–16017.

27. Clausen, M., et al., High-force generation is a conserved property of type IV pilus systems. Journal of Bacteriology, 2009. 191(14): p. 4633–4638.

28. Denisin, A.K. and B.L. Pruitt, Tuning the range of polyacrylamide gel stiffness for mechanobiology applications. ACS applied materials & interfaces, 2016. 8(34): p. 21893–21902.

29. Normand, V., et al., New insight into agarose gel mechanical properties. Biomacromolecules, 2000. 1(4): p. 730–738.

30. Chui, M.M., R.J. Phillips, and M.J. McCarthy, Measurement of the porous microstructure of hydrogels by nuclear magnetic resonance. Journal of colloid and interface science, 1995. 174(2): p. 336–344.

31. Holmes, D.L. and N.C. Stellwagen, Estimation of polyacrylamide gel pore size from Ferguson plots of linear DNA fragments. II. Comparison of gels with different crosslinker concentrations, added agarose and added linear polyacrylamide. Electrophoresis, 1991. 12(9): p. 612–619.

32. Jiang, L. and S. Granick, Real-space, in situ maps of hydrogel pores. ACS nano, 2016. 11(1): p. 204–212.

33. Narayanan, J., J.-Y. Xiong, and X.-Y. Liu. Determination of agarose gel pore size: Absorbance measurements vis a vis other techniques. in Journal of Physics: Conference Series. 2006. IOP Publishing.

34. Koch, M.D., et al., Competitive binding of independent extension and retraction motors explains the quantitative dynamics of type IV pili. Proceedings of the National Academy of Sciences, 2021. 118(8).

35. Konopka, M.C., et al., Crowding and confinement effects on protein diffusion in vivo. Journal of Bacteriology, 2006. 188(17): p. 6115–6123.

36. Kumar, M., M.S. Mommer, and V. Sourjik, Mobility of cytoplasmic, membrane, and DNA-binding proteins in Escherichia coli. Biophysical Journal, 2010. 98(4): p. 552–559.

37. Kilmury, S.L. and L.L. Burrows, Type IV pilins regulate their own expression via direct intramembrane interactions with the sensor kinase PilS. Proceedings of the National Academy of Sciences, 2016. 113(21): p. 6017–6022.

38. Biais, N., et al., Force-dependent polymorphism in type IV pili reveals hidden epitopes. Proceedings of the National Academy of Sciences, 2010. 107(25): p. 11358–11363.

39. Merz, A.J., M. So, and M.P. Sheetz, Pilus retraction powers bacterial twitching motility. Nature, 2000. 407(6800): p. 98–102.

40. Ellison, C.K., et al., Retraction of DNA-bound type IV competence pili initiates DNA uptake during natural transformation in Vibrio cholerae. Nature microbiology, 2018: p. 1.

41. Ribbe, J., et al., The role of cyclic di-GMP and exopolysaccharide in type IV pilus dynamics. Journal of Bacteriology, 2017: p. JB. 00859–16.

42. Skerker, J.M. and H.C. Berg, Direct observation of extension and retraction of type IV pili. Proceedings of the National Academy of Sciences, 2001. 98(12): p. 6901–6904.

43. Chlebek, J.L., et al., Fresh extension of Vibrio cholerae competence type IV pili predisposes them for motor-independent retraction. Applied and Environmental Microbiology, 2021: p. AEM. 00478–21.

44. Beaussart, A., et al., Nanoscale adhesion forces of Pseudomonas aeruginosa type IV pili. ACS nano, 2014. 8(10): p. 10723–10733.

45. Touhami, A., et al., Nanoscale characterization and determination of adhesion forces of Pseudomonas aeruginosa pili by using atomic force microscopy. Journal of Bacteriology, 2006. 188(2): p. 370–377.

46. Lu, S., et al., Nanoscale pulling of type IV pili reveals their flexibility and adhesion to surfaces over extended lengths of the pili. Biophysical Journal, 2015. 108(12): p. 2865–2875.

47. Coggan, K.A. and M.C. Wolfgang, Global regulatory pathways and cross-talk control Pseudomonas aeruginosa environmental lifestyle and virulence phenotype. Current issues in molecular biology, 2011. 14(1): p. 47–70.

48. Guimarães, C.F., et al., The stiffness of living tissues and its implications for tissue engineering. Nature Reviews Materials, 2020. 5(5): p. 351–370.

49. Inclan, Y.F., et al., A scaffold protein connects type IV pili with the Chp chemosensory system to mediate activation of virulence signaling in Pseudomonas aeruginosa. Molecular Microbiology, 2016. 101(4): p. 590–605.

50. Sourjik, V. and N.S. Wingreen, Responding to chemical gradients: bacterial chemotaxis. Current opinion in cell biology, 2012. 24(2): p. 262–268.

51. Del Medico, L., et al., The type IV pilin PilA couples surface attachment and cell-cycle initiation in Caulobacter crescentus. Proceedings of the National Academy of Sciences, 2020. 117(17): p. 9546–9553.

52. Snyder, R.A., et al., Surface sensing stimulates cellular differentiation in Caulobacter crescentus. Proceedings of the National Academy of Sciences, 2020. 117(30): p. 17984–17991.

53. Wozniak, M.A. and C.S. Chen, Mechanotransduction in development: a growing role for contractility. Nature Reviews Molecular Cell Biology, 2009. 10(1): p. 34–43.

54. Ortega, D.R., et al., Assigning chemoreceptors to chemosensory pathways in Pseudomonas aeruginosa. Proceedings of the National Academy of Sciences, 2017. 114(48): p. 12809–12814.

55. Hmelo, L.R., et al., Precision-engineering the Pseudomonas aeruginosa genome with two-step allelic exchange. Nature protocols, 2015. 10(11): p. 1820.

56. Koch, M.D. and J.W. Shaevitz, Introduction to Optical Tweezers, in Optical Tweezers - Methods and Protocols, A. Gennerich, Editor 2016, Springer: New York.

57. Herrick, W.G., et al., PEG-phosphorylcholine hydrogels as tunable and versatile platforms for mechanobiology. Biomacromolecules, 2013. 14(7): p. 2294–2304.

58. Subhash, G., et al., Concentration dependence of tensile behavior in agarose gel using digital image correlation. Experimental Mechanics, 2011. 51(2): p. 255–262.

59. Tuson, H.H., et al., Measuring the stiffness of bacterial cells from growth rates in hydrogels of tunable elasticity. Molecular Microbiology, 2012. 84(5): p. 874–891.

60. Craig, L., M.E. Pique, and J.A. Tainer, Type IV pilus structure and bacterial pathogenicity. Nature Reviews Microbiology, 2004. 2(5): p. 363.

61. Bustamante, C., et al., Mechanical processes in biochemistry. Annual review of biochemistry, 2004. 73(1): p. 705–748.

62. Grassia, P., E. Hinch, and L. Nitsche, Computer simulations of Brownian motion of complex systems. Journal of Fluid Mechanics, 1995. 282: p. 373–403.

63. Bertrand, J.J., J.T. West, and J.N. Engel, Genetic analysis of the regulation of type IV pilus function by the Chp chemosensory system of Pseudomonas aeruginosa. Journal of Bacteriology, 2010. 192(4): p. 994–1010.

64. Whitchurch, C.B., et al., Pseudomonas aeruginosa fimL regulates multiple virulence functions by intersecting with Vfr-modulated pathways. Molecular Microbiology, 2005. 55(5): p. 1357–1378.

